# Development of a Pan-Proteome Multi-Epitope Vaccine Candidate Against Nipah Virus Using Reverse Vaccinology and Immunoinformatics Approaches

**DOI:** 10.64898/2026.11.05.764467

**Authors:** Rasel Khan Nishan, Md. Nahian, Bithi Akter, Tabassum Basher, Tanjum Ahmed Nodee, Irin Aminur Asha, Mostafizur Rahaman Sany, Sanjida Akter Sathy, Farzana Sharmin Shila, Mohammad Ariful Islam, Abul Kalam Azad, Md. Shariful Islam

## Abstract

Nipah virus (NiV) is a high-consequence zoonotic pathogen for which no licensed human vaccine is currently available. This study applied a consensus-guided pan-proteome immunoinformatics framework to design and prioritize a multi-epitope vaccine candidate using sequence information from all nine NiV proteins. A total of 901 NiV protein sequences were retrieved, of which 842 passed quality filtering and were processed through clustering, multiple-sequence alignment, and protein-specific consensus generation. Consensus-derived fragments were screened for cytotoxic T-lymphocyte, helper T-lymphocyte, and B-cell epitopes, followed by multi-criterion filtering based on antigenicity, allergenicity, toxicity, conservancy, class-specific immune-response properties, and human-proteome similarity. Ten epitope-selection strategies were combined with two adjuvant candidates and three construct architectures to generate 60 vaccine constructs. Shortlisted candidates were subsequently evaluated for physicochemical properties, HLA population coverage, structural characteristics, conformational B-cell epitopes, TLR3 docking, molecular dynamics, immune-response simulation, and expression-oriented optimization. VC033_V6_BD_L03 construct was prioritized as the lead candidate and showed combined predicted population coverage of 99.78% worldwide and 99.96% in South Asia. Docking against human TLR3 structure 1ZIW yielded a ClusPro score of −1624.8. During a 100-ns molecular-dynamics simulation, the complex underwent early conformational reorganization followed by lower late-phase structural variation and persistent interchain hydrogen bonding. MM/GBSA analysis yielded a mean interaction-energy estimate of −20.52 ± 15.42 kcal/mol. C-ImmSim predicted coordinated humoral and cellular responses, while codon optimization generated a 1,539-bp expression sequence compatible with virtual cloning into pET-28a(+). These analyses prioritize VC033_V6_BD_L03 as a consensus-derived, pan-proteome NiV multi-epitope vaccine candidate for experimental evaluation of expression, immunogenicity, safety, and protective efficacy.

## Introduction

Nipah virus (NiV) is a highly pathogenic zoonotic henipavirus that causes severe respiratory and neurological disease in humans. The virus was first identified during the 1998–1999 outbreak in Malaysia and Singapore and has since caused recurrent outbreaks, particularly in Bangladesh and India [1, 2]. Fruit bats of the genus Pteropus constitute the principal natural reservoir, and human infection can result from contact with infected animals, consumption of contaminated food products, or direct person-to-person transmission. NiV infection is associated with an estimated case-fatality rate of approximately 40–75%, although this varies among outbreaks and healthcare settings. Recent outbreaks in Bangladesh and India continue to demonstrate the persistence of the threat, and NiV remains a priority pathogen under the World Health Organization R&D Blueprint. No licensed vaccine or NiV-specific therapeutic is currently available, although several vaccine platforms are progressing through preclinical and clinical development [3, 4]. The ChAdOx1 NipahB candidate entered the first Phase II NiV vaccine trial in Bangladesh in 2025, representing an important advance in clinical development while underscoring the continuing need for complementary vaccine strategies [5].

NiV is an enveloped, non-segmented, negative-sense RNA virus whose six transcriptional units encode nine proteins: nucleocapsid protein (N), phosphoprotein (P), matrix protein (M), fusion protein (F), attachment glycoprotein (G), large polymerase protein (L), and the accessory proteins V, W, and C. The surface glycoproteins G and F mediate receptor attachment and membrane fusion and are major targets of neutralizing-antibody responses. N, P, and L form essential components of the viral replication machinery, while M coordinates virion assembly and budding. The P-gene products P, V, W, and C also contribute to antagonism of host antiviral signaling, particularly interferon-mediated responses [2, 6, 7]. The distinct biological roles and sequence characteristics of these proteins provide a broader antigenic repertoire than strategies restricted to the major surface glycoproteins alone.

Most advanced NiV vaccine platforms have focused on the F and/or G glycoproteins because of their central roles in viral entry and their ability to induce neutralizing antibodies. Nevertheless, protective antiviral immunity also involves cellular responses against infected cells and coordinated CD4 T-cell help in addition to antibody-mediated recognition. A multi-epitope approach offers the possibility of combining B-cell epitopes with MHC class I-restricted cytotoxic T-lymphocyte (CTL) and MHC class II-restricted helper T-lymphocyte (HTL) epitopes within a defined construct. Such an approach can also prioritize conserved regions and broaden representation across HLA alleles and viral proteins. Reverse vaccinology and immunoinformatics provide a practical framework for this type of screening by enabling large sequence datasets to be evaluated computationally before experimental candidate testing [8, 9]. This is particularly useful for pathogens such as NiV, for which experimental work with infectious virus requires high-containment facilities.

A growing number of computational studies have applied these approaches to NiV vaccine design, although the source proteins and selection strategies have varied considerably. Soltan et al. (2021) [10] initially evaluated the NiV proteome and subsequently selected epitopes from F, G, V, and W for multi-epitope vaccine construction. Kumar et al. (2024) [11] focused on glycoprotein G and nucleocapsid N, whereas Sharma et al. (2025) [12] screened the six structural proteins G, F, M, N, P, and L. Broader proteome-based strategies have also been reported. Srivastava et al. (2023) [13] predicted CTL, HTL, and B-cell epitopes from all nine NiV proteins, while Banico et al. (2024) [14] incorporated nine NiV proteins and used multiple sequence alignments followed by Protein Variability Server analysis to identify highly conserved sequence fragments. Thus, use of the complete NiV protein repertoire itself is not unprecedented; rather, previous studies differ in how sequence diversity is represented before epitope prediction and in how candidate epitopes are subsequently assembled and prioritized.

Sequence variation among circulating NiV isolates creates an additional consideration for epitope-based design because candidates derived from a single reference sequence may not optimally represent the diversity present across available viral sequences. Consensus-based antigen strategies offer one means of reducing dependence on an individual isolate while preserving commonly represented sequence features. This concept has already been explored experimentally for NiV surface antigens: consensus F- and G-based DNA vaccines have been evaluated for immunogenicity, and a consensus F antigen has demonstrated protective efficacy against representative NiV strains in an animal model [15, 16]. However, these approaches have centered on one or two glycoproteins rather than applying protein-level consensus generation across the complete nine-protein repertoire. Similarly, the previously reported all-protein immunoinformatics studies did not use the same workflow of protein-specific sequence clustering, consensus generation, ambiguity resolution, and subsequent systematic multi-architecture construct screening used here.

Accordingly, the present study applied a consensus-guided pan-proteome reverse-vaccinology and immunoinformatics framework for the design and prioritization of a NiV multi-epitope vaccine candidate. Sequence datasets representing G, F, N, M, L, P, V, W, and C were quality filtered and clustered before protein-specific consensus sequences were generated and ambiguity-free conserved fragments were obtained for epitope prediction. Predicted CTL, HTL, and B-cell epitopes were subjected to sequential screening based on antigenicity, allergenicity, toxicity, sequence conservancy, class-specific immune-response characteristics, and human-proteome similarity. Rather than assembling a single predefined vaccine sequence, multiple epitope-selection strategies, adjuvant candidates, and sequence architectures were systematically evaluated to identify favorable construct combinations. Shortlisted constructs were subsequently assessed through HLA population-coverage analysis, secondary- and tertiary-structure prediction, structural refinement, conformational B-cell epitope analysis, receptor docking, molecular-dynamics simulation, immune-response simulation, and expression-oriented optimization. An overview of the complete computational workflow is presented in Figure 1. This integrated strategy was designed to prioritize a conserved and broadly HLA-representative NiV multi-epitope candidate for subsequent experimental evaluation.

**Figure 1.**
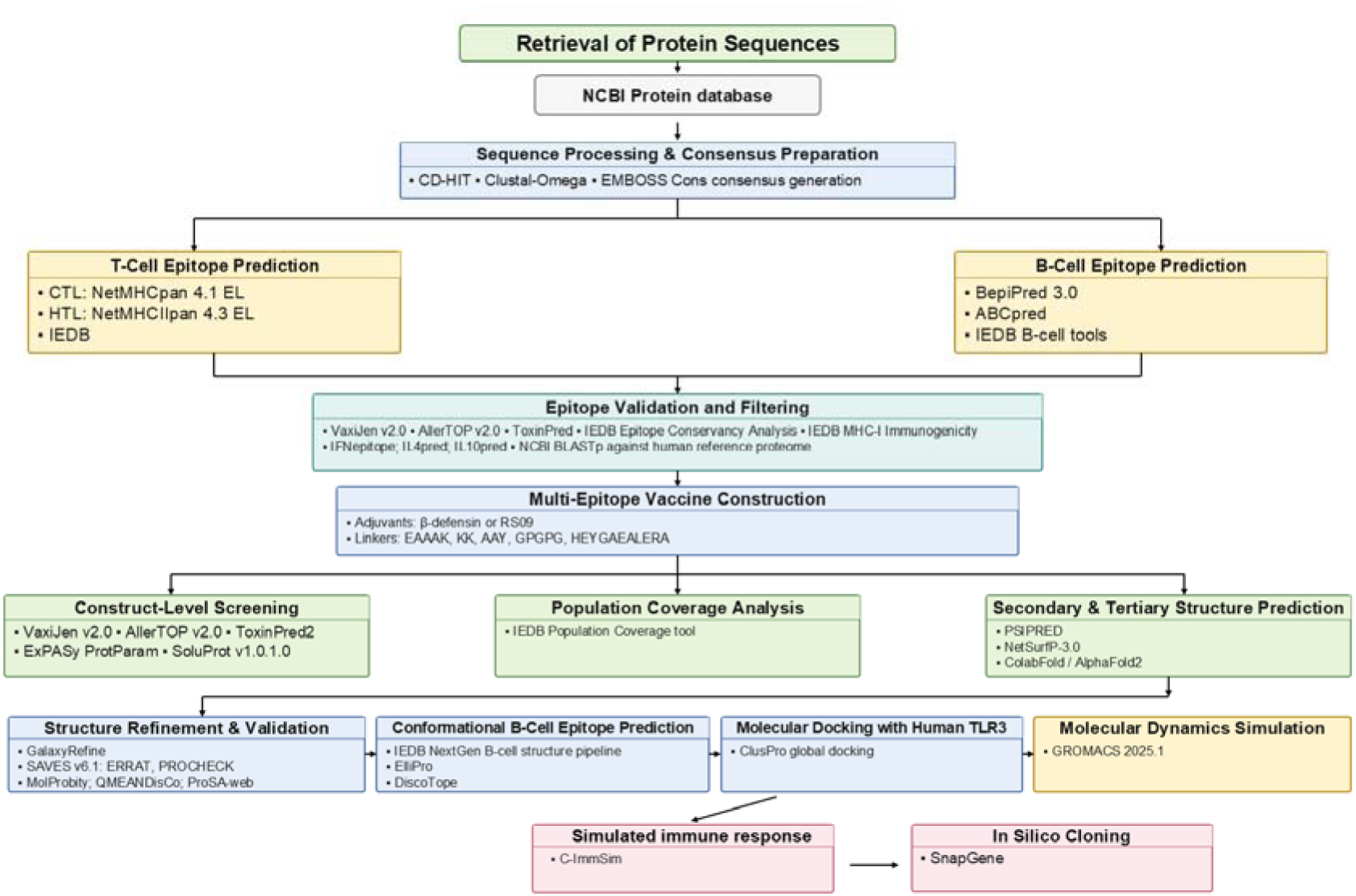
Overall workflow for the pan-proteome reverse vaccinology and immunoinformatics-based design of a multi-epitope vaccine candidate against Nipah henipavirus.

## Methods and Materials

### Sequence data collection and consensus preparation

Complete amino acid sequences of nine major Nipah henipavirus proteins—glycoprotein (G), fusion protein (F), nucleocapsid protein (N), matrix protein (M), large polymerase protein (L), phosphoprotein (P), and the accessory proteins V, W, and C—were retrieved from the NCBI Protein database using protein-specific search queries (Supplementary file 1). Each protein dataset was processed independently to facilitate pan-proteome epitope screening.

The retrieved FASTA datasets were quality-filtered, and sequences containing ambiguous or non-standard amino acid residues were excluded. To minimize sequence redundancy, the filtered datasets were independently clustered using CD-HIT [17] at a 99% sequence identity threshold, with minimum alignment coverage of 99% for both the shorter and longer sequences, a word size of 5, and global sequence identity calculation.

Representative sequences from each CD-HIT cluster were subjected to multiple sequence alignment using Clustal Omega [18]. For proteins represented by a single non-redundant sequence, multiple sequence alignment was omitted. Protein-specific consensus sequences were subsequently generated from the alignments using EMBOSS Cons [19]. Gap characters were removed from the resulting consensus sequences, and positions containing ambiguous residues (X) were treated as breakpoints to generate conserved, ambiguity-free sequence fragments. These fragments were subsequently used for epitope prediction. Fragments with a minimum length of 9 and 15 amino acids were retained for cytotoxic T-lymphocyte (CTL) and helper T-lymphocyte (HTL) epitope prediction, respectively.

### Epitope prediction

For cytotoxic T-lymphocyte (CTL) epitope prediction, MHC class I ligand predictions were generated using the Next-Generation Immune Epitope Database (IEDB) T Cell Prediction, Class I (TC1) server and the DTU NetMHCpan 4.1 server [20, 21]. The human 27-allele MHC-I reference panel from IEDB server was used to provide broad HLA representation (Supplementary File 2). In the IEDB TC1 workflow, the NetMHCpan 4.1 eluted-ligand (EL) prediction method was used with the server-default peptide-length settings, whereas the DTU NetMHCpan 4.1 server was run using its default EL-prediction settings. Prediction records were evaluated using the EL percentile rank (%Rank_EL), and peptides with %Rank_EL ≤1.0 were retained for downstream analysis.

Helper T-lymphocyte (HTL) epitopes were predicted using the Next-Generation IEDB T Cell Prediction, Class II (TC2) server and the DTU NetMHCIIpan 4.3 server [20, 22]. Predictions were performed against a human 27-allele MHC-II from IEDB reference panel (Supplementary File 2). The IEDB TC2 workflow used the NetMHCIIpan EL prediction method with the default peptide-processing settings, while NetMHCIIpan 4.3 was independently run through the DTU web server using its default EL-prediction settings. HTL prediction records with %Rank_EL ≤10.0 were retained for subsequent analysis.

Linear B-cell epitope prediction was performed using BepiPred 3.0 and ABCpred. BepiPred 3.0 predictions were obtained through both the IEDB B-cell sequence-prediction workflow and the DTU BepiPred 3.0 server [23]. In the IEDB workflow, BepiPred Linear Epitope Prediction 3.0 was run using the Majority Vote and Variable Threshold options with the default advanced settings. The DTU BepiPred 3.0 server was independently run using its default settings. These two BepiPred runs represented separate server implementations of the same underlying prediction method and were therefore treated as supporting prediction evidence rather than as independent algorithms. As an algorithmically distinct predictor, ABCpred was applied using a prediction threshold of 0.51, a fixed window length of 16 amino acids, and the overlapping-prediction option enabled [24]. Duplicate predictions arising from overlapping segments were removed. B-cell epitope candidates shorter than six amino acids were excluded.

### Epitope validation, filtering, and Human-proteome similarity screening

Predicted epitopes retained after class-specific prediction filtering were subjected to sequential validation for antigenicity, allergenicity, toxicity, sequence conservancy, and epitope-class-specific immune-response potential. Antigenicity was assessed using VaxiJen v2.0 with the target organism set to virus and a threshold of 0.4 [25]. Only epitopes classified as probable antigens were retained. Allergenicity was evaluated using AllerTOP v2.0 with the default server settings, and only epitopes classified as probable non-allergens were retained [26]. Toxicity was assessed using ToxinPred with the default server settings, and peptides classified as non-toxic were retained [27]. Sequence conservancy was evaluated using the IEDB Epitope Conservancy Analysis tool against the corresponding CD-HIT-derived non-redundant representative sequences of each source protein [28]. Epitopes exhibiting >95% sequence conservancy were retained for further analysis. CTL epitopes were additionally evaluated using the IEDB MHC-I Immunogenicity tool, which estimates the immunogenic potential of MHC-I-presented peptides based on sequence characteristics [29]. Only CTL epitopes with an immunogenicity score >0 were retained. HTL epitopes were evaluated for their predicted ability to induce interferon-gamma (IFN-γ), interleukin-4 (IL-4), and interleukin-10 (IL-10) using the human-host IFNepitope2 server, IL4pred, and IL10pred, respectively [30, 31, 32]. These tools were run using their default server settings. Only HTL epitopes classified as inducers by all three cytokine-prediction tools were retained.

Following these validation steps, the retained CTL, HTL, and B-cell epitopes were screened for sequence similarity to human proteins using NCBI BLASTp against the RefSeq Protein database, restricted to Homo sapiens (taxid: 9606) [33]. The automatic short-query adjustment applied by the NCBI BLASTp web interface was retained. A peptide was considered to have a full-length exact human match only when the alignment exhibited 100% sequence identity, the alignment length equaled the complete peptide length, and the alignment spanned the query from the first to the final residue. Epitopes meeting these criteria were excluded from subsequent vaccine-construct design.

### Vaccine construct design and construct-level screening

Following epitope validation, 10 different epitope sets (V1–V10) were generated to explore alternative vaccine designs. Each set contained 27 epitopes: nine B-cell, nine CTL, and nine HTL epitopes. The selection strategies prioritized different features, including antigenicity, CTL immunogenicity, HLA-binding breadth, cytokine-inducing potential, sequence conservancy, low similarity to human proteins, and diversity of epitope positions. Exact duplicate peptide sequences were avoided within each epitope set. The detailed selection criteria for V1–V10 are provided in Supplementary file 3. Each epitope set was combined with one of two N-terminal adjuvants: human β-defensin-3 (hBD-3) or the synthetic TLR4 agonist peptide RS09 [34, 35]. The adjuvant was joined to the epitope region using an EAAAK linker. Three construct arrangements were evaluated. In L01, CTL, HTL, and B-cell epitopes were arranged in separate blocks without linkers between individual epitopes. In L02 and L03, CTL, HTL, and B-cell epitopes were separated by AAY, GPGPG, and KK linkers, respectively. The major epitope blocks were separated by HEYGAEALERAG. L02 used the order B-cell–CTL–HTL, whereas L03 used CTL–HTL–B-cell. Combining 10 epitope sets, two adjuvants, and three construct arrangements produced 60 vaccine constructs.

All 60 constructs were evaluated for antigenicity using VaxiJen v2.0, allergenicity using AllerTOP v2.0, toxicity using ToxinPred2, physicochemical properties using ExPASy ProtParam, and predicted soluble expression using SoluProt [25, 26, 36, 37, 38]. Constructs were retained for structural analysis if they were predicted to be antigenic, non-allergenic, non-toxic, and soluble, with a VaxiJen score ≥0.90, SoluProt score ≥0.70, instability index ≤35, GRAVY ≤−0.10, and aliphatic index ≥80.

### Population coverage analysis

Population coverage analysis was performed using the IEDB Population Coverage tool (https://tools.iedb.org/population/) to estimate the expected HLA-based coverage of selected vaccine constructs [39]. Because this analysis is based on MHC-restricted T-cell responses, only CTL and HTL epitopes and their corresponding HLA restrictions were included; B-cell epitopes were excluded from the population coverage calculation.

For each selected construct, CTL epitopes were paired with their predicted HLA-I binding alleles, and HTL epitopes were paired with their predicted HLA-II binding alleles. Duplicate epitope–allele pairs were removed before submission. Three analyses were performed for each construct: MHC-I coverage using CTL epitopes, MHC-II coverage using HTL epitopes, and combined MHC-I plus MHC-II coverage using both T-cell epitope classes. Coverage was estimated for the World population and the South Asia population to evaluate both global relevance and regional relevance to areas where NiV outbreaks have repeatedly occurred.

### Secondary- and tertiary-structure prediction

Secondary structure of the selected vaccine constructs was predicted using PSIPRED and NetSurfP-3.0 [40, 41]. PSIPRED was used to estimate the proportions of α-helices, β-strands, and coils, while NetSurfP-3.0 provided an independent secondary-structure prediction together with relative solvent accessibility and disorder estimates. PSIPRED and NetSurfP-3.0 are established sequence-based tools for secondary-structure and related structural-feature prediction. Three-dimensional structures were predicted using the ColabFold implementation of AlphaFold2 [42, 43]. Predictions were performed in monomer mode using MMseqs2-based multiple-sequence alignments, with templates disabled, five models, three recycles, and no AMBER relaxation. The rank-1 ColabFold model was selected for subsequent refinement, and mean pLDDT was used as a measure of prediction confidence. ColabFold combines AlphaFold2 structure prediction with accelerated MMseqs2-based sequence searches.

### Structure refinement and validation

The rank-1 ColabFold model of each construct was refined using the GalaxyRefine web server with the default settings [44]. GalaxyRefine improves protein models through side-chain rebuilding and structural relaxation. The refined models were evaluated using PROCHECK and ERRAT through the SAVES server, together with MolProbity, QMEANDisCo, and ProSA-web [45, 46, 47, 48, 49]. PROCHECK was used to assess Ramachandran statistics, ERRAT to evaluate overall structural quality, MolProbity to assess all-atom geometry and steric clashes, QMEANDisCo to estimate model quality, and ProSA-web to obtain the Z-score. These tools provide complementary assessments of stereochemistry, atomic contacts, and overall model quality.

### Conformational B-cell epitope prediction

Conformational B-cell epitopes were predicted from the selected refined vaccine structures using the IEDB NextGen structure-based B-cell epitope prediction pipeline. Before submission, each selected GalaxyRefine model was standardized by retaining heavy atoms, assigning the complete vaccine structure to chain A, and renumbering residues sequentially. Where available, residue-level pLDDT values from the corresponding rank-1 ColabFold model were transferred to the B-factor field of the standardized PDB file. ElliPro was used as the primary predictor of discontinuous B-cell epitopes, with the maximum clustering distance set to 6 Å [50]. DiscoTope-3.0 was applied to the same structures as a residue-level cross-check [51]. Both analyses used the standardized chain A structures submitted as AlphaFold-type models through the IEDB pipeline. ElliPro discontinuous epitopes were treated as the primary conformational B-cell epitope predictions. For each ElliPro epitope, overlap with residues classified as epitope residues by DiscoTope-3.0 was calculated and used as supporting evidence.

### Molecular docking with human TLR3

Molecular docking was performed to evaluate predicted interactions between the selected vaccine constructs and human Toll-like receptor 3 (TLR3). Two human TLR3 extracellular-domain structures, PDB IDs 1ZIW and 2A0Z, were used as receptors. Receptor structures were prepared by retaining the protein chain and removing non-protein components, including water molecules, ions, glycans, and other HETATM records. Refined vaccine structures were similarly prepared as protein docking partners. The 10 refined vaccine constructs were initially subjected to protein–protein docking against both TLR3 structures using ClusPro in global docking mode without predefined interaction restraints [52], generating 20 receptor–vaccine combinations. Docking solutions were evaluated according to the ClusPro ranking and scoring scheme. Following ClusPro screening, the prioritized VC033_V6_BD_L03–1ZIW pair was independently evaluated using the HDOCK server [53]. The prepared TLR3 and vaccine protein structures were submitted as receptor and ligand, respectively, using the default HDOCK protein–protein docking workflow. The top-ranked HDOCK-generated complex from VC033_V6_BD_L03–1ZIW was selected as the starting structure for subsequent molecular dynamics simulation and was used as returned by the docking workflow.

### Molecular Dynamics Simulation

Molecular dynamics simulation of the selected TLR3–VC033 complex was performed using GROMACS 2025.1 [54]. Both components were treated as proteins. The supplied production topology was generated using gmx pdb2gmx with the CHARMM27 force field [55] and TIP3P water model [56]. The system was subjected to steepest-descent energy minimization for 2,500 steps, followed by 100 ps NVT and 100 ps NPT equilibration with position restraints applied to protein heavy atoms.

The production simulation was performed for 100 ns using a 2 fs integration time step at 300 K and 1 bar. Temperature was controlled using the V-rescale thermostat [57] and pressure using the Parrinello–Rahman barostat [58]. Periodic boundary conditions were applied in all three dimensions, long-range electrostatic interactions were treated using the particle-mesh Ewald method [59], and bond constraints were applied using LINCS [60]. Real-space Coulomb and van der Waals cutoffs were 1.0 nm. Compressed trajectory coordinates were written every 5,000 steps, corresponding to 10 ps intervals. The production topology contained the two protein chains and 145,177 TIP3P water molecules; no explicit ion molecules were listed in the supplied production topology, and the GROMACS production log reported a total system charge of +23.

Structural analyses were performed using GROMACS trajectory-analysis tools [54]. Backbone RMSD was calculated after least-squares fitting to the Backbone group. Residue-wise RMSF was evaluated from Cα atoms; the supplied RMSF output corresponded to TLR3 residues 30–664. Radius of gyration and solvent-accessible surface area were calculated using the GROMACS Protein group, with a 0.14-nm solvent probe used for SASA analysis. Intermolecular hydrogen bonds were evaluated between chain A and chain B using a donor– acceptor distance cutoff of 0.35 nm and a donor–hydrogen–acceptor angular criterion of ≥120°.

MM/GBSA calculations were performed using gmx_MMPBSA v1.6.3 [61]. A total of 1,001 snapshots were sampled uniformly from the 100-ns production trajectory. The Generalized Born calculation used igb = 5, corresponding to the GB-OBC2 model, while the nonpolar solvation contribution was estimated using the LCPO approach. Binding-energy components included van der Waals, electrostatic, polar-solvation, and nonpolar-solvation terms. Entropic contributions were not included.

Principal component analysis (PCA) was performed from Cα positional fluctuations after structural alignment [62]. The principal components were obtained from the covariance matrix, and trajectory projections were examined along the dominant components. The free-energy landscape (FEL) was constructed from the PC1–PC2 conformational distribution using the Boltzmann relation. Dynamic cross-correlation matrix (DCCM) analysis was used to examine correlated and anticorrelated Cα motions during the trajectory [63].

### In silico immune simulation

The predicted immune response to the prioritized vaccine construct VC033_V6_BD_L03 was evaluated using the C-ImmSim online server [64]. The complete 513-amino-acid construct sequence was used as the antigen. The primary simulation comprised two administrations on days 0 and 28, corresponding to C-ImmSim time steps 1 and 84. A secondary exploratory simulation incorporated a third administration on day 56 (time step 168), resulting in a three-dose schedule of days 0, 28, and 56. Each simulation was run for 1,050 C-ImmSim time steps, corresponding to approximately 350 days. The HLA configuration comprised HLA-A03:01, HLA-A11:01, HLA-B15:01, HLA-B35:01, HLA-DRB103:01, and HLA-DRB104:01. The random seed was set to 12345, and the server volume parameter was 10. Each administration was submitted as “vaccine (no LPS)”, with an antigen quantity parameter of 1,000 and a C-ImmSim adjuvant parameter of 100. The same VC033_V6_BD_L03 sequence was used for each administration. Immune-response outputs were evaluated over the complete simulated period, including B-cell and helper T-cell populations and their memory compartments, cytotoxic T-cell responses, immunoglobulin profiles, antigen kinetics, and cytokine/interleukin responses. The two-dose simulation was used as the primary analysis, while the three-dose simulation was used to examine the predicted response to an additional booster administration.

### In Silico Expression Optimization and Virtual Cloning

The 513-amino-acid VC033_V6_BD_L03 construct was back-translated into a 1,539-bp coding sequence using EMBOSS Backtranseq with the Escherichia coli K-12 codon-usage table [19]. The resulting nucleotide sequence was subsequently optimized for E. coli expression using the GenSmart Codon Optimization Tool [65]. Codon-usage properties of the optimized sequence were assessed using the GenRCA Rare Codon Analysis Tool, including the codon adaptation index (CAI), codon frequency distribution (CFD), and GC content [66]. The optimized coding sequence was further analyzed using RNAfold to predict mRNA secondary structure and minimum free energy [67]. Before virtual cloning, the sequence was screened for internal NdeI and BamHI restriction sites. NdeI and BamHI sites were then incorporated at the 5′ and 3′ ends of the construct, respectively, with a stop codon positioned before the BamHI site. The resulting expression cassette was virtually inserted between the corresponding restriction sites of the pET-28a(+) expression vector using SnapGene [68].

## Results

### Data collection and consensus preparation

A total of 901 Nipah virus protein sequences representing nine viral proteins were retrieved from NCBI. After quality filtering, 842 sequences were retained and 59 sequences containing non-standard amino acid residues were excluded. The retained dataset included 105 G, 113 F, 125 N, 107 M, 31 L, 104 P, 85 V, 82 W, and 90 C protein sequences. CD-HIT clustering reduced the 842 sequences to 57 non-redundant representative sequences: 4 G, 3 F, 2 N, 2 M, 2 L, 10 P, 12 V, 14 W, and 8 C sequences. Consensus-sequence analysis generated 11 ambiguity-free fragments for downstream epitope prediction. G, F, N, M, L, P, and C each produced a single consensus sequence, whereas ambiguous positions in the V and W consensus sequences resulted in two conserved fragments for each protein.

### Epitope prediction

Epitope prediction across the 11 consensus-derived fragments generated 659,869 CTL, 118,706 HTL, and 886 B-cell prediction records. After applying the predefined prediction thresholds, 10,521 CTL, 12,624 HTL, and 696 B-cell records were retained for further screening. Because the same T-cell peptide could be predicted for more than one HLA allele, the retained CTL and HTL results contained repeated peptide sequences. After exact sequence deduplication, 3,318 unique CTL and 3,410 unique HTL epitopes remained. The B-cell dataset contained 696 unique candidate epitopes after processing. These non-redundant epitope sets were subsequently subjected to antigenicity, allergenicity, toxicity, conservancy, and class-specific validation.

### Epitope Validation, Filtering, and Human-proteome similarity screening

The predicted epitopes were subjected to sequential validation based on antigenicity, allergenicity, toxicity, sequence conservancy, and epitope-class-specific immune-response criteria. Application of the predefined stringent filtering criteria substantially reduced the candidate pool, yielding 384 CTL, 59 HTL, and 127 B-cell epitopes. Overall, 570 epitopes satisfied all required validation criteria and were retained as the final candidate pool for human-proteome similarity screening. The 570 validated epitopes were subsequently screened against the *Homo sapiens* reference proteome using BLASTp to identify peptides exhibiting complete sequence identity with human proteins. None of the CTL, HTL, or B-cell epitopes showed a full-length exact match (100% identity across the complete peptide length) to a human protein. Consequently, no candidates were excluded at this stage, and all 570 validated epitopes were retained for subsequent multi-epitope vaccine construct design (Supplementary file 4).

### Vaccine construction and construct-level validation

Ten epitope sets (V1–V10) were generated, each containing 27 non-redundant epitopes comprising nine CTL, nine HTL, and nine B-cell epitopes. Eight epitope sets represented all nine NiV proteins, whereas V4 and V8 represented seven and six proteins, respectively. Combination of the 10 epitope sets with two adjuvants and three construct architectures generated 60 vaccine constructs, with sequence lengths ranging from 389 to 517 amino acids. No construct contained an epitope showing a full-length exact match to a human protein. Construct-level screening showed that all 60 constructs were predicted to be antigenic and non-toxic, while 58 were classified as non-allergenic and 58 as stable. SoluProt classified 30 constructs as soluble (Supplementary file 5). Subsequent prioritization identified 11 constructs for structural analysis. The 11 selected constructs originated from epitope sets V1, V4, V5, V6, and V10, and all contained the hBD-3 adjuvant. Five used the L02 architecture, five used L03, and one used L01. Their VaxiJen scores ranged from 0.9063 to 1.1559 and SoluProt scores from 0.7223 to 0.7742. Instability indices ranged from 28.08 to 33.58, GRAVY values from −0.291 to −0.157, and aliphatic indices from 84.89 to 96.88. The selected constructs were 433–516 amino acids long, with predicted molecular weights of approximately 49.6–57.0 kDa (Table 2 and Supplementary file 6). Among the shortlisted constructs, VC033_V6_BD_L03 was carried forward through the downstream structural and receptor-interaction analyses and was ultimately prioritized as the final vaccine candidate. The organization of the 513-amino-acid construct is shown in Figure 2.

**Table 1.** Principal characteristics of the 11 vaccine constructs retained after construct-level screening. All selected constructs contained the human β-defensin-3 (hBD-3) adjuvant and were predicted to be non-allergenic, non-toxic, stable, and soluble. V1–V10 denote the epitope-selection strategies, whereas L01–L03 denote the construct architectures. MW, molecular weight; GRAVY, grand average of hydropathicity.

| **Construct** | **VaxiJen score** | **SoluProt score** | **Instability index** | **GRAVY** | **Aliphatic index** |
|---|---|---|---|---|---|
| VC002_V1_BD_L02 | 1.0914 | 0.7253 | 33.58 | –0.257 | 90.74 |
| VC003_V1_BD_L03 | 1.0667 | 0.7388 | 33.17 | –0.257 | 90.74 |
| VC020_V4_BD_L02 | 1.1559 | 0.7713 | 29.92 | –0.169 | 94.34 |
| VC021_V4_BD_L03 | 1.1465 | 0.7742 | 29.92 | –0.169 | 94.34 |
| VC025_V5_BD_L01 | 0.9585 | 0.7480 | 31.17 | –0.162 | 96.88 |
| VC026_V5_BD_L02 | 0.9787 | 0.7260 | 28.73 | –0.291 | 84.89 |
| VC027_V5_BD_L03 | 0.9656 | 0.7342 | 28.08 | –0.291 | 84.89 |
| VC032_V6_BD_L02 | 1.0533 | 0.7495 | 32.58 | –0.201 | 91.54 |
| **VC033_V6_BD_L03** | **1.0321** | **0.7607** | **32.17** | **–0.201** | **91.54** |
| VC056_V10_BD_L02 | 0.9093 | 0.7225 | 33.30 | –0.157 | 93.84 |
| VC057_V10_BD_L03 | 0.9063 | 0.7223 | 33.45 | –0.157 | 93.84 |

**Figure 2.**
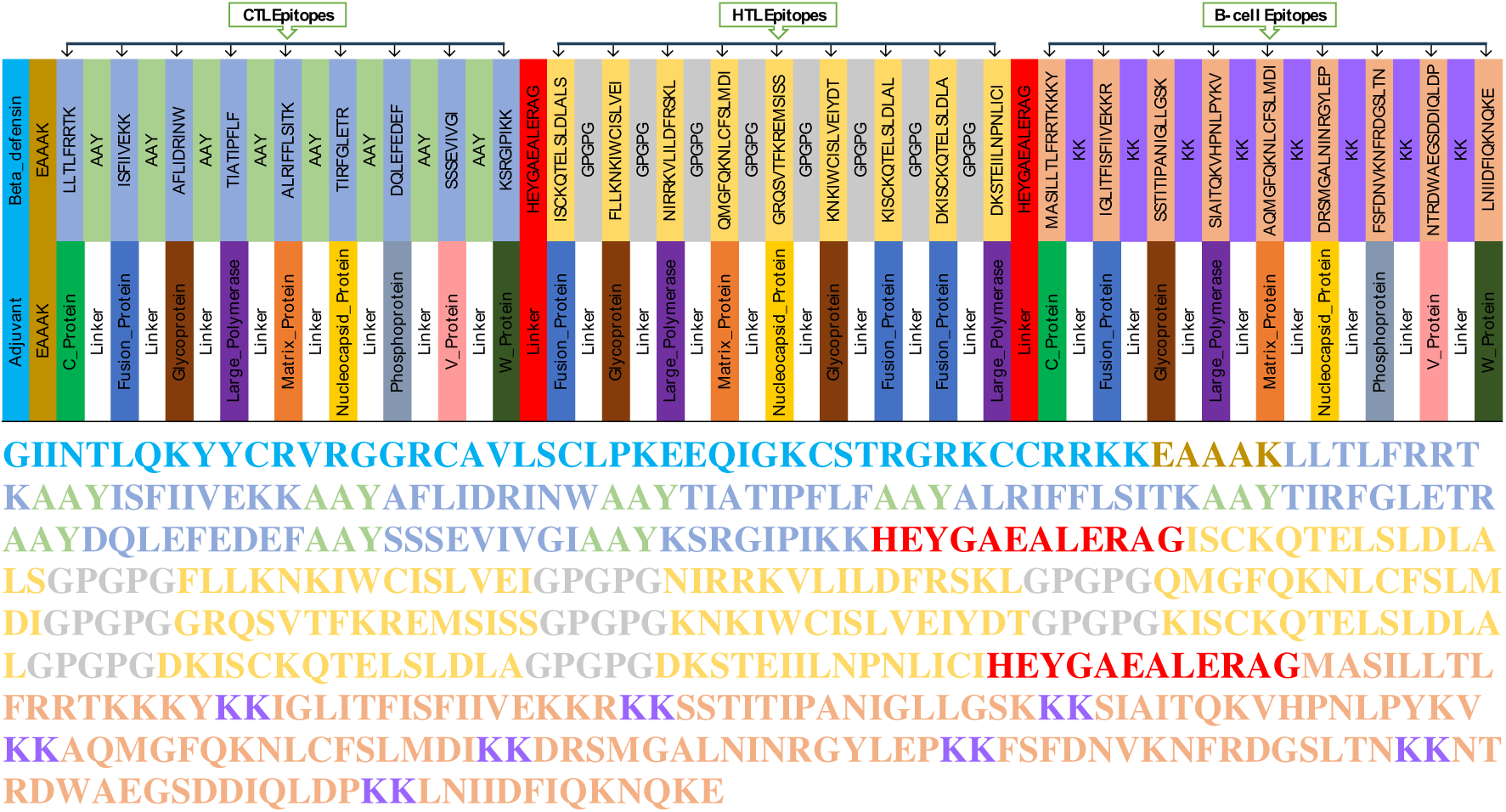
Schematic organization of the prioritized multi-epitope vaccine construct VC033_V6_BD_L03. The 513-amino-acid construct contains an N-terminal human β-defensin-3 (hBD-3) adjuvant followed by nine CTL, nine HTL, and nine B-cell epitopes arranged according to the L03 architecture. The adjuvant is connected to the epitope region by an EAAAK linker. CTL, HTL, and B-cell epitopes are separated by AAY, GPGPG, and KK linkers, respectively, while HEYGAEALERAG linkers separate the three major epitope blocks.

**Table 2.** ClusPro molecular docking results of selected vaccine constructs with receptors 1ZIW and 2A0Z. ClusPro docking results showing cluster size, center score, and lowest energy score for each construct–receptor complex. More negative scores indicate more favorable predicted docking.

| Version | Receptor | Cluster_members | Center_score | Lowest_energy_score |
|---|---|---|---|---|
| VC002_L02 | 1ZIW | 62 | -1281.6 | -1456.4 |
| VC003_L03 | 1ZIW | 68 | -1108.8 | -1178.8 |
| VC020_L02 | 1ZIW | 104 | -1051.9 | -1094.9 |
| VC021_L03 | 1ZIW | 33 | -1238 | -1238 |
| VC026_L02 | 1ZIW | 45 | -1385.2 | -1488.4 |
| VC027_L03 | 1ZIW | 46 | -980.5 | -1229.6 |
| VC032_L02 | 1ZIW | 34 | -1155.8 | -1321 |
| **VC033_L03** | 1ZIW | 60 | -1515.8 | -1624.8 |
| VC056_L02 | 1ZIW | 38 | -1068.8 | -1466.9 |
| VC057_L03 | 1ZIW | 38 | -1221.5 | -1539 |
| VC002_L02 | 2A0Z | 46 | -1029.3 | -1164.8 |
| VC003_L03 | 2A0Z | 41 | -1070.5 | -1216.6 |
| VC020_L02 | 2A0Z | 55 | -1057.6 | -1099.6 |
| VC021_L03 | 2A0Z | 72 | -1165.7 | -1256.8 |
| VC026_L02 | 2A0Z | 39 | -1307.8 | -1414.9 |
| VC027_L03 | 2A0Z | 59 | -990.5 | -1125.5 |
| VC032_L02 | 2A0Z | 26 | -1337.8 | -1393.1 |
| VC033_L03 | 2A0Z | 50 | -989.7 | -1285.3 |
| VC056_L02 | 2A0Z | 35 | -1088.8 | -1103.4 |
| VC057_L03 | 2A0Z | 147 | -277.2 | -337.7 |

**(A)** **(B)** **(C)**

### Population coverage analysis

The 11 shortlisted vaccine constructs showed consistently high predicted HLA population coverage. Combined MHC-I + MHC-II coverage ranged from 99.73–99.97% worldwide and 99.92–100% in South Asia. The highest combined coverage was observed for VC025_V5_BD_L01, VC026_V5_BD_L02, and VC027_V5_BD_L03, each reaching 99.97% worldwide and 100% in South Asia. The subsequently prioritized construct VC033_V6_BD_L03 showed a predicted combined population coverage of 99.78% worldwide and 99.96% in South Asia (Supplementary file 7). Its coverage profiles indicated that the majority of individuals were predicted to recognize multiple epitope–HLA combinations, with cumulative coverage approaching 100% in both populations (Figure 3A,B).

**Figure 3.**
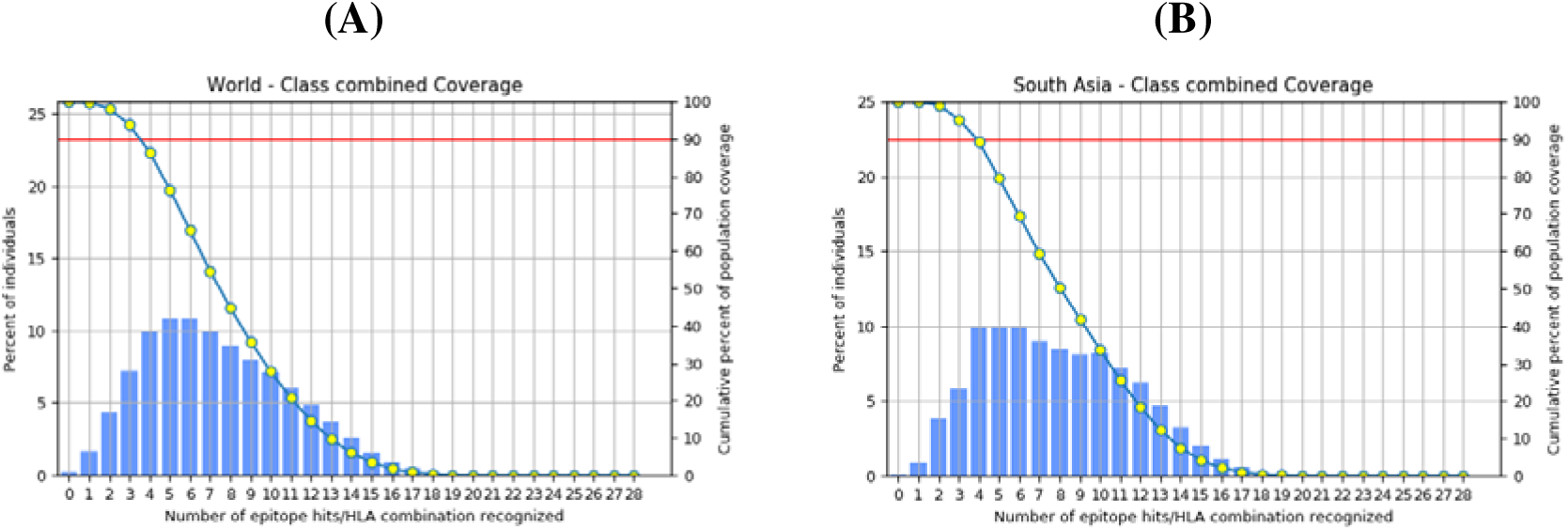
Predicted combined MHC-I and MHC-II population coverage of VC033_V6_BD_L03. Population-coverage profiles of the prioritized VC033_V6_BD_L03 construct for (A) the World population and (B) the South Asian population, calculated using the IEDB Population Coverage tool. Blue bars represent the percentage of individuals predicted to recognize a given number of epitope–HLA combinations, and the line represents the cumulative percentage of population coverage. The red horizontal line indicates 90% population coverage. Combined predicted coverage reached 99.78% worldwide and 99.96% in South Asia.

### Prediction of the secondary and tertiary structure of constructed vaccine

Secondary-structure predictions were successfully obtained for all 11 shortlisted vaccine constructs. PSIPRED predicted 20.71–35.08% α-helical, 22.40–33.53% β-strand, and 40.55– 47.34% coil content across the constructs. NetSurfP-3.0 similarly indicated substantial helical and coil content, with predicted α-helices ranging from 31.64–47.17%, β-strands from 4.88– 12.02%, and coils from 45.03–58.40% (Supplementary file 8). For the subsequently prioritized VC033_V6_BD_L03, PSIPRED predicted 30.02% α-helix, 29.43% β-strand, and 40.55% coil, whereas NetSurfP-3.0 predicted 34.70% α-helix, 8.97% β-strand, and 56.34% coil. NetSurfP-3.0 additionally yielded a mean relative solvent accessibility of 0.313 and a mean disorder score of 0.0914 for this construct. The residue-level secondary-structure profiles of VC033_V6_BD_L03 are shown in **Figure 4A,B**. ColabFold generated five tertiary-structure models for each of the 11 constructs, and a rank-1 model was obtained for every candidate. Mean pLDDT values of the rank-1 models ranged from 30.99 to 39.36, with VC033_V6_BD_L03 showing a mean pLDDT of 39.12 (Supplementary file 9). These rank-1 models were subsequently carried forward for structural refinement and validation.

**Figure 4.**
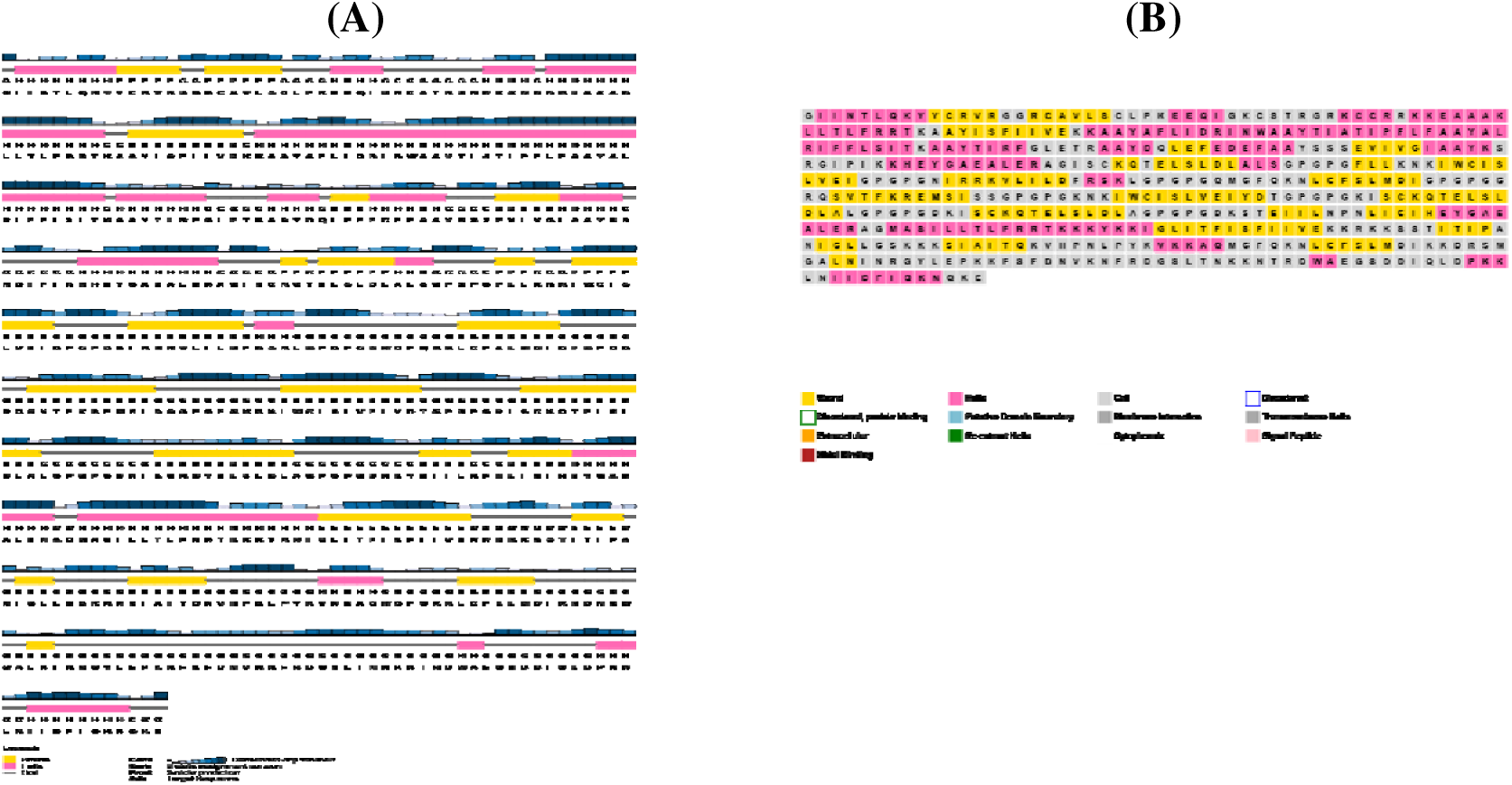
Predicted secondary-structure profile of the prioritized vaccine construct VC033_V6_BD_L03. (A) PSIPRED secondary-structure prediction across the 513-amino-acid construct, showing residue-level assignments of α-helical, β-strand, and coil regions together with prediction confidence. (B) NetSurfP-3.0 residue-level structural-feature representation of VC033_V6_BD_L03. PSIPRED predicted 30.02% α-helix, 29.43% β-strand, and 40.55% coil, whereas NetSurfP-3.0 predicted 34.70% α-helix, 8.97% β-strand, and 56.34% coil.

### Vaccine construct refinement and validation

All 11 shortlisted vaccine constructs were subjected to GalaxyRefine, generating five refined models per construct. Following structural-quality assessment, 10 constructs yielded a refined model that was retained for downstream analysis. Across these selected models, Ramachandran favored-region percentages ranged from 93.6–95.2%, ERRAT overall quality factors from 80.14–96.33, MolProbity scores from 0.84–1.16, ProSA Z-scores from −3.51 to approximately 0, and QMEANDisCo average local scores from 0.2445–0.3196. Complete structural-validation results for the selected refined models are provided in Supplementary file 10. For VC033_V6_BD_L03, GalaxyRefine model 1 was retained. PROCHECK analysis showed 93.9% of residues in favored regions, 5.4% in additionally allowed regions, and 0.7% in disallowed regions, while ERRAT produced an overall quality factor of 91.192 (Figure 5A,B). Standalone MolProbity analysis yielded a MolProbity score of 0.99, a clashscore of 0.61, 0.00% poor rotamers, 95.89% Ramachandran-favored residues, and 0.59% Ramachandran outliers. The ProSA Z-score was −1.79, and the QMEANDisCo average local score was 0.295 ± 0.053. QMEAN4 and QMEAN6 Z-scores were −5.47 and −5.77, respectively. GalaxyRefine reported a GDT-HA of 0.8601, an RMSD of 0.721 Å, and a GalaxyRefine MolProbity score of 1.163 for the selected model.

**Figure 5.**
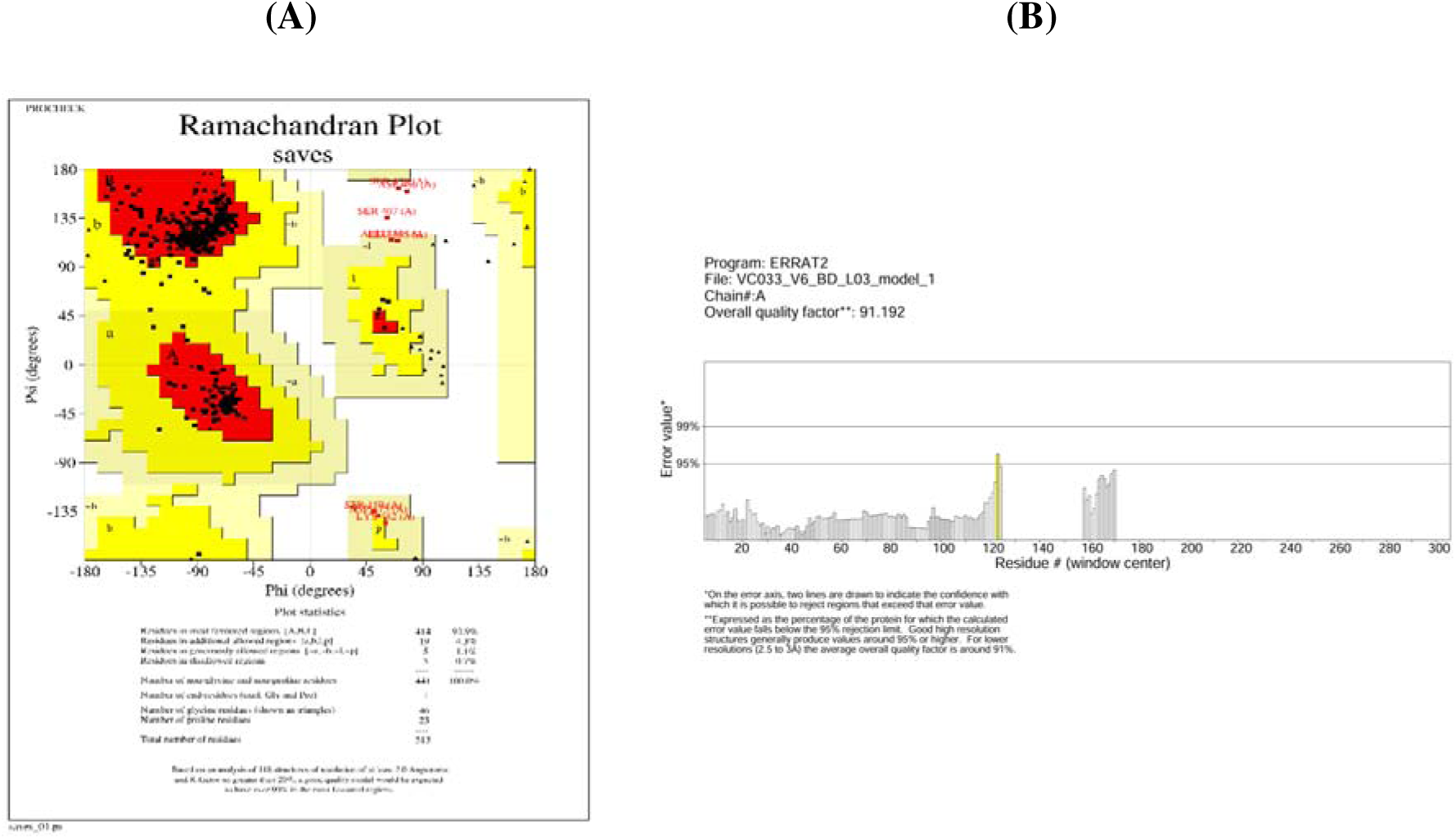
Structural validation of the refined VC033_V6_BD_L03 model. (A) PROCHECK Ramachandran plot of the selected GalaxyRefine model, showing the distribution of residues across favored, additionally allowed, generously allowed, and disallowed conformational regions. (B) ERRAT analysis of the same refined model, showing residue-wise error values and an overall quality factor of 91.192.

### Screening of conformational B-cell epitopes

Structure-based B-cell epitope prediction was completed for the 10 selected refined vaccine models. ElliPro identified 235 discontinuous epitopes, with 20–29 epitopes per construct and scores ranging from 0.503 to 0.980. ElliPro additionally identified 87 linear epitopes, corresponding to 5–12 epitopes per construct. DiscoTope-3.0 identified 65 epitope stretches, with 5–9 stretches per construct. For VC033_V6_BD_L03, ElliPro predicted 26 discontinuous epitopes with scores ranging from 0.509 to 0.946, together with five linear epitopes. DiscoTope-3.0 identified seven epitope stretches in the same refined model. Of the 26 ElliPro discontinuous epitopes, 20 showed residue-level overlap with DiscoTope-3.0 epitope assignments. Overall, 75 of 233 residues (32.2%) comprising the ElliPro discontinuous epitopes overlapped DiscoTope-positive residues, and 10 epitopes showed ≥50% residue overlap. Detailed epitope-level results are provided in Supplementary file 11.

### Molecular docking analysis

The 10 selected refined vaccine constructs were docked against two TLR3 structures, 1ZIW and 2A0Z, generating 20 receptor–construct combinations. For TLR3 structure 1ZIW, VC033_V6_BD_L03 produced the lowest model-0 energy score (−1624.8), followed by VC057_V10_BD_L03 (−1539.0) and VC026_V5_BD_L02 (−1488.4). Their corresponding model-0 center scores were −1515.8, −1221.5, and −1385.2, respectively. The three complexes are shown in Figure 6A–C. Complete docking scores for all receptor–construct combinations are provided in Table 2. Among the complexes generated with 2A0Z, VC026_V5_BD_L02 produced the lowest model-0 energy score (−1414.9), followed by VC032_V6_BD_L02 (−1393.1) and VC033_V6_BD_L03 (−1285.3). Based on the ClusPro ranking, the 1ZIW–VC033_V6_BD_L03 complex was carried forward for additional docking with HDOCK. HDOCK model 1 returned a docking score of −314.68 and a confidence score of 0.9642, and this top-ranked complex was selected for subsequent molecular-dynamics simulation.

**Figure 6.**
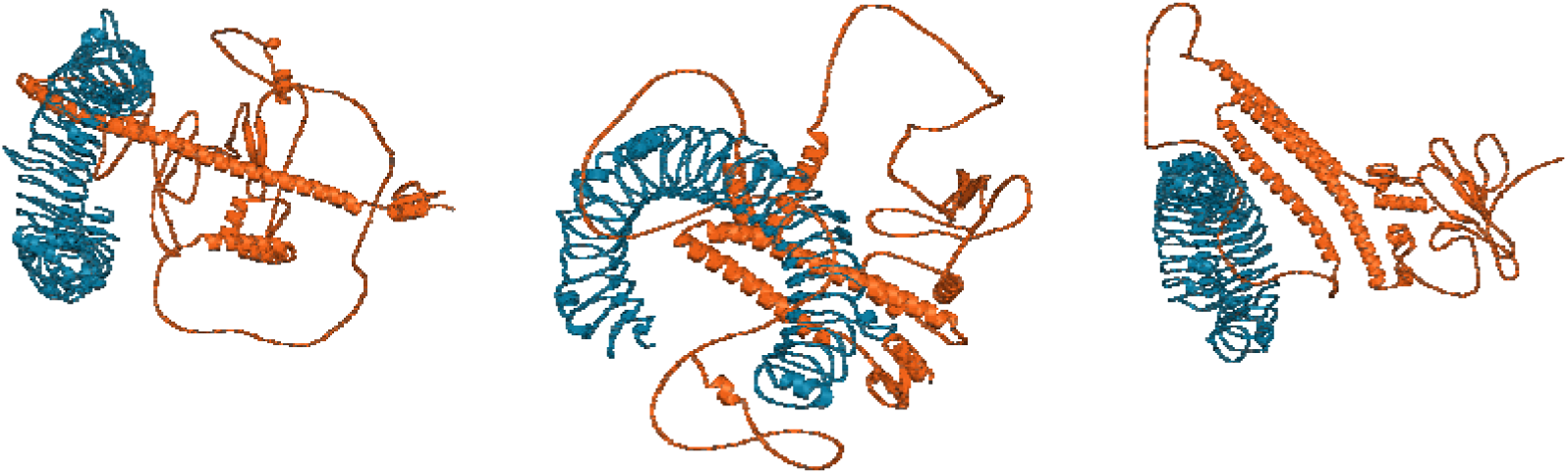
Top-ranked ClusPro docking complexes of selected vaccine constructs with TLR3 structure 1ZIW. Structural representations of the three vaccine constructs with the lowest model-0 ClusPro energy scores when docked to TLR3 (PDB ID: 1ZIW): (A) VC033_V6_BD_L03, score −1624.8; (B) VC057_V10_BD_L03, score −1539.0; and (C) VC026_V5_BD_L02, score −1488.4. TLR3 is shown in blue and the vaccine constructs in orange.

### Molecular Dynamics Simulation

The TLR3–VC033_V6_BD_L03 complex showed substantial structural adjustment during the early phase of the 100-ns trajectory. Backbone RMSD increased rapidly after the start of production and continued to rise more gradually until approximately 35–40 ns, after which the trajectory fluctuated within a narrower range. During the final 50 ns, the mean RMSD was 1.250 ± 0.033 nm, with a range of 1.131–1.366 nm (Figure 7A). The Protein-group radius of gyration decreased during the early trajectory and subsequently showed reduced variation. During 50–100 ns, the mean Rg was 3.431 ± 0.017 nm, ranging from 3.378 to 3.489 nm (Figure 7B). Residue-wise Cα RMSF analysis of TLR3 residues 30–664 yielded a mean fluctuation of 0.190 ± 0.090 nm. The highest fluctuation was observed at residue 662 (0.764 nm), while most internal residues displayed lower fluctuations than terminal regions (Figure 7C). Intermolecular hydrogen-bond analysis showed relatively few contacts during the first approximately 15 ns, followed by more frequent interchain interactions during the remainder of the simulation. During the final 50 ns, the complex formed an average of 2.67 ± 1.14 intermolecular hydrogen bonds per frame, with a maximum of seven, and at least one interchain hydrogen bond was present in approximately 98.4% of sampled frames (Figure 7D). Collectively, the RMSD, Rg, RMSF, and hydrogen-bond profiles showed an early conformational reorganization followed by a more stable late-phase conformational regime with persistent receptor–construct contacts.

**Figure 7.**
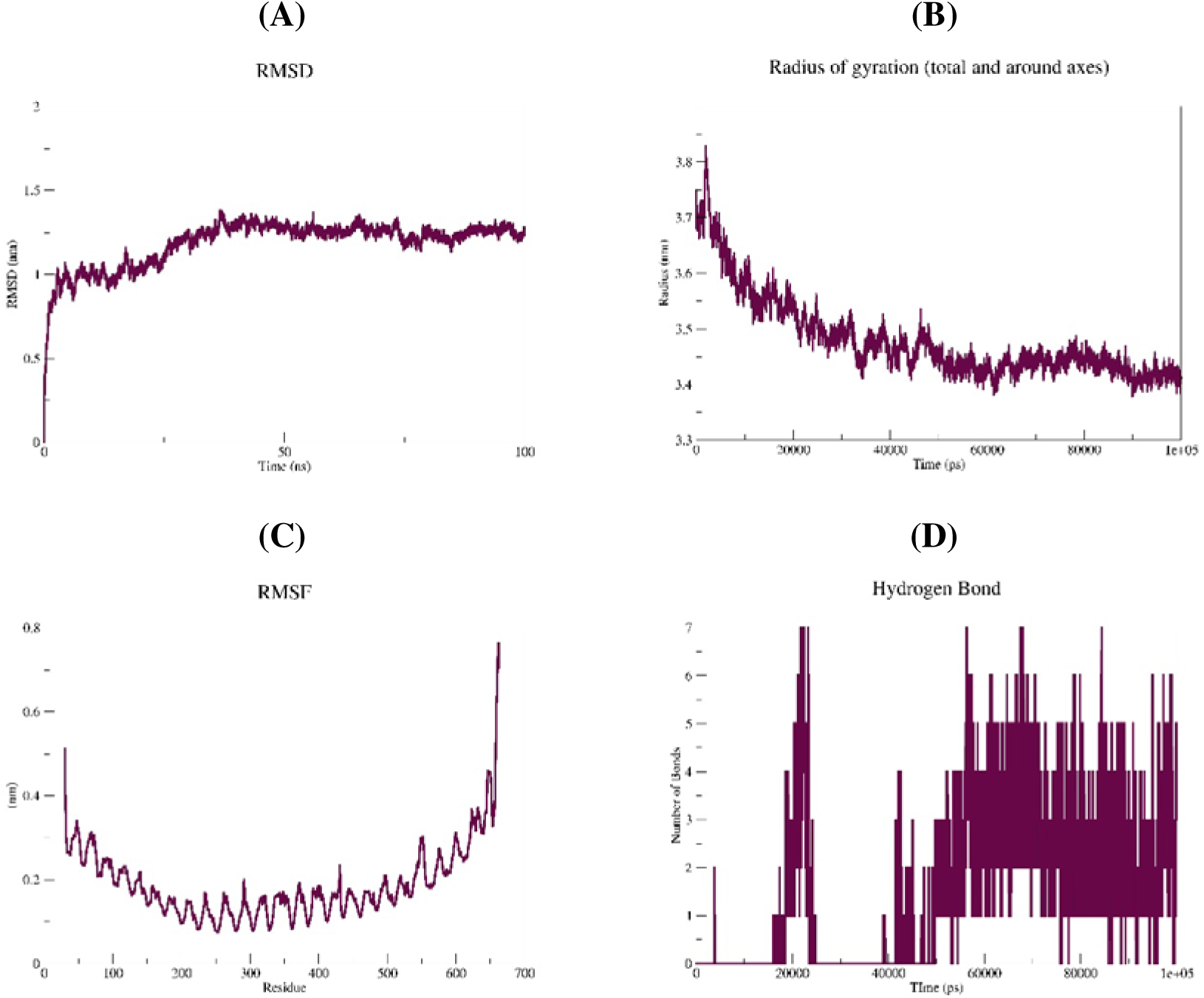
Molecular-dynamics analysis of the TLR3–VC033_V6_BD_L03 complex during the 100-ns production trajectory. (A) Backbone root-mean-square deviation (RMSD). (B) Protein-group radius of gyration (Rg). (C) Cα root-mean-square fluctuation (RMSF) of TLR3 residues 30–664. (D) Number of intermolecular hydrogen bonds between the TLR3 and vaccine-construct chains throughout the trajectory.

The Protein-group solvent-accessible surface area averaged 532.77 ± 2.67 nm² during the final 50 ns and showed comparatively limited variation over this interval (Supplementary file 12 Figure 1A). Principal component analysis showed that PC1 and PC2 accounted for 38.66% and 29.37% of the total positional variance, respectively, corresponding to 68.03% cumulatively. PC3 contributed a further 7.52%, while the first six principal components together accounted for 86.2% of the total variance (Figure 8). Trajectory projections along PC1–PC2, PC2–PC3, and PC1–PC3 occupied multiple regions of principal-component space (Supplementary file 12 Figure 1B–D). The PC1–PC2 free-energy landscape contained multiple low-energy basins within an approximately 0–10 kJ/mol range, representing several energetically accessible conformational states sampled during the trajectory (Supplementary file 12 Figure 1E). DCCM analysis showed both positively and negatively correlated Cα motions across the analyzed coordinates (Supplementary file 12 Figure 1F). Residue-specific DCCM assignments were not made because a residue-to-matrix-index mapping was not available in the supplied output.

**Figure 8.**
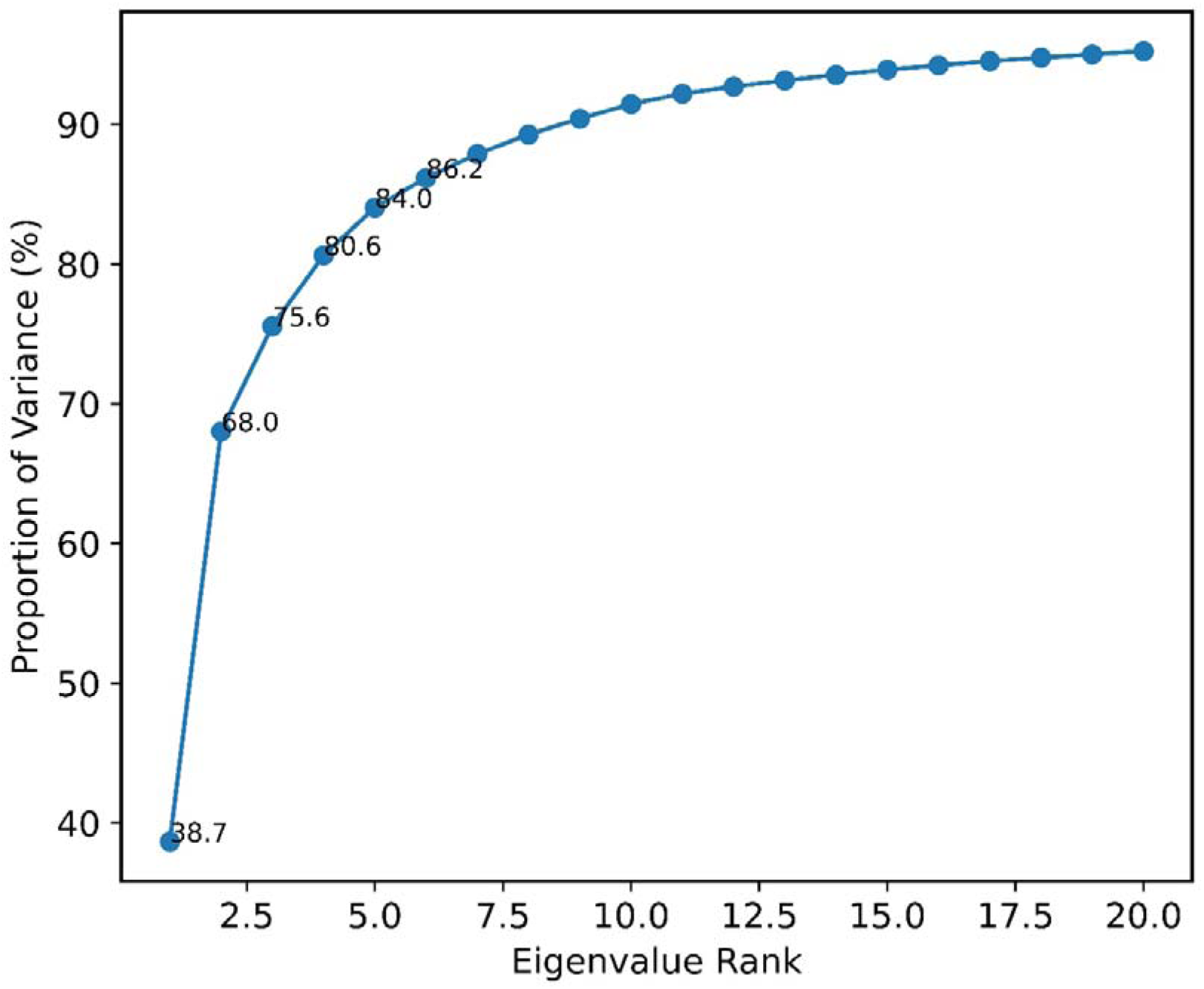
Cumulative variance explained by PCA. The first principal component accounts for 38.7% of the total variance, while the first two, three, four, five, and six components cumulatively explain 68.0%, 75.6%, 80.6%, 84.0%, and 86.2%, respectively. The cumulative variance exceeds 90% within approximately the first 9–10 principal components and gradually approaches around 95% by the 20th component.

MM/GBSA analysis of 1,001 trajectory snapshots yielded a mean total binding-energy estimate of −20.52 ± 15.42 kcal/mol. The van der Waals and electrostatic contributions were −24.22 ± 18.07 kcal/mol and −113.35 ± 69.78 kcal/mol, respectively, whereas the polar-solvation contribution was +121.12 ± 74.63 kcal/mol and the nonpolar-solvation contribution was −4.07 ± 3.02 kcal/mol (Table 3). The negative total MM/GBSA energy was driven primarily by favorable electrostatic and van der Waals contributions, which outweighed the opposing polar-solvation term and supported an energetically favorable receptor–construct association across the sampled trajectory conformations.

**Table 3.** MM/GBSA energy components of the TLR3–VC033_V6_BD_L03 complex calculated from 1,001 snapshots of the 100-ns production trajectory (mean ± SD, kcal/mol).

| **Energy component** | **Mean ± SD (kcal/mol)** |
|---|---|
| VDWAALS | −24.22 ± 18.07 |
| EEL | −113.35 ± 69.78 |
| EGB | +121.12 ± 74.63 |
| ESURF | −4.07 ± 3.02 |
| GGAS | −137.57 ± 86.27 |
| GSOLV | +117.05 ± 71.83 |
| TOTAL | −20.52 ± 15.42 |

### Simulated immune response

The primary two-dose C-ImmSim simulation predicted coordinated humoral and cellular responses following administration of VC033_V6_BD_L03 on days 0 and 28. B-cell populations increased following immunization, with memory B-cell populations remaining detectable throughout the simulated period. Helper T-cell populations also increased during the early response, and a persistent helper T-cell memory population was maintained through the end of the approximately 350-day simulation (Figure 9A,B). The cytotoxic T-cell compartment showed a transient increase in activated cells following immunization, followed by a progressive shift toward the resting population during the later simulation period (Figure 9C). Additional cellular outputs, including plasma B cells, natural killer cells, macrophages, dendritic cells, and epithelial-cell populations, are provided in Supplementary File 13. The humoral response showed a marked increase following the second administration. Antigen levels declined after each exposure, while the booster was followed by a substantially greater combined IgM + IgG response, together with IgM, IgG1, and IgG2 components. Antibody levels subsequently declined from their peak but remained detectable throughout the simulated period (Figure 9D). Cytokine analysis showed discrete responses following vaccine administration, with prominent IFN-γ and IL-2 peaks associated with the immunization periods. Additional cytokines and interleukins, including IL-4, IL-12, TGF-β, TNF-α, IL-10, IL-6, IL-18, IL-8, and IL-23, were also represented in the simulated response (Figure 9E). The exploratory three-dose simulation produced an additional immune-response peak following the day-56 administration. A third antigen exposure was accompanied by renewed B-cell and helper T-cell responses, an additional rise in immunoglobulin production, and a third cytokine response. Memory B-cell and helper T-cell populations remained detectable through the end of the simulation. The complete three-dose response profiles are presented in Supplementary File 14.

**Figure 9.**
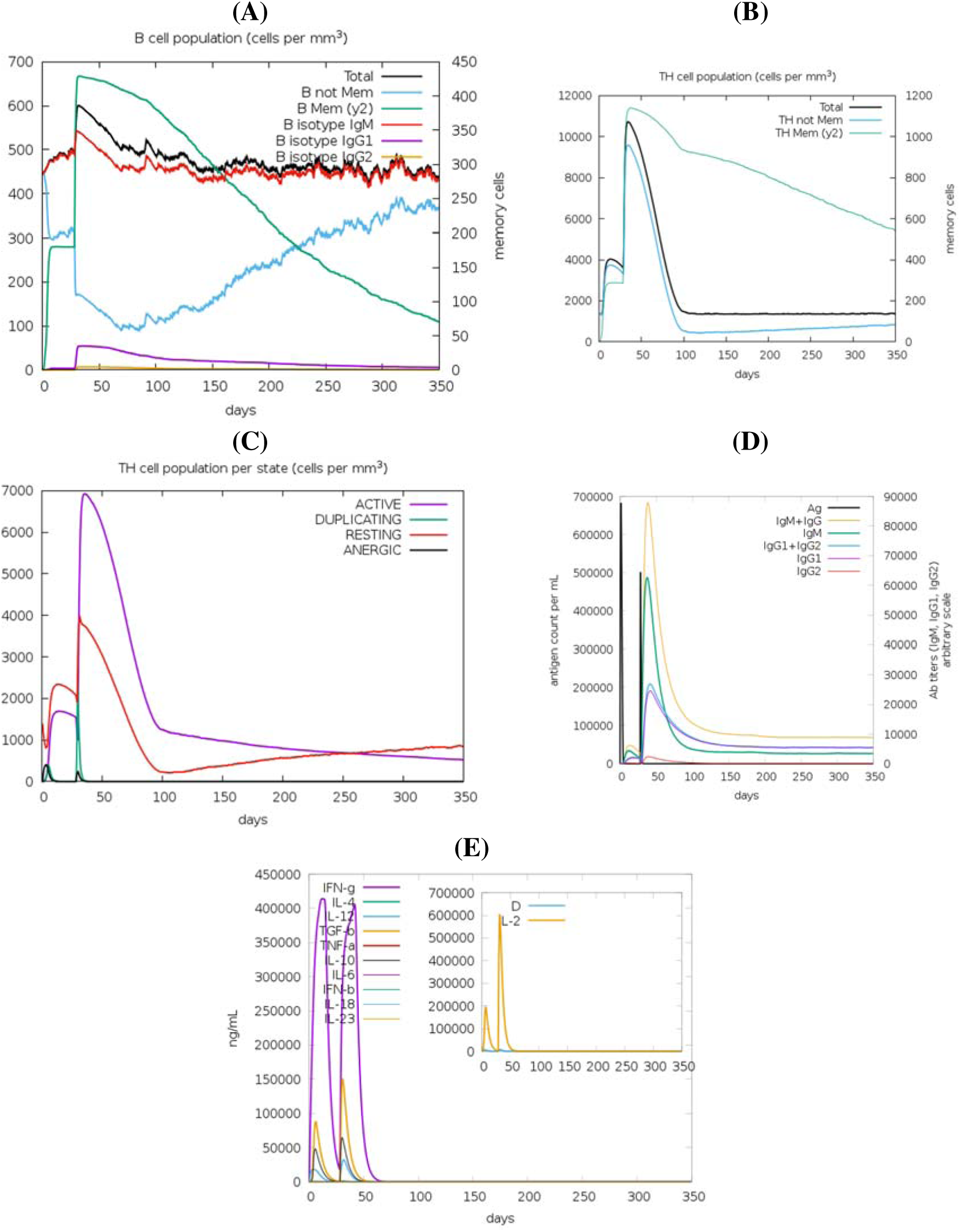
C-ImmSim-predicted immune response to the two-dose VC033_V6_BD_L03 vaccination schedule. The construct was administered on simulated days 0 and 28. **(A)** B-cell population and memory-cell dynamics. **(B)** Helper T-cell population and memory-cell dynamics. **(C)** Cytotoxic T-cell population according to activation state. **(D)** Antigen and immunoglobulin kinetics, including combined IgM + IgG, IgM, IgG1, and IgG2 responses. **(E)** Predicted cytokine and interleukin concentrations during the simulation, including IFN-γ and IL-2 responses. The simulation was performed for approximately 350 days using C-ImmSim.

### In Silico Expression Optimization and Virtual Cloning

The optimized VC033_V6_BD_L03 coding sequence retained a length of 1,539 bp and showed a CAI of 0.75, CFD of 0.04, and GC content of approximately 50.3%. RNAfold predicted a minimum free energy of −497.40 kcal/mol for the optimized mRNA sequence. Restriction-site analysis identified no internal NdeI or BamHI sites, permitting directional insertion of the optimized coding sequence between these restriction sites in the pET-28a(+) vector. Virtual cloning in SnapGene generated an in silico recombinant plasmid of 6,878 bp containing the optimized VC033_V6_BD_L03 coding sequence (**Figure 10**).

**Figure 10.**
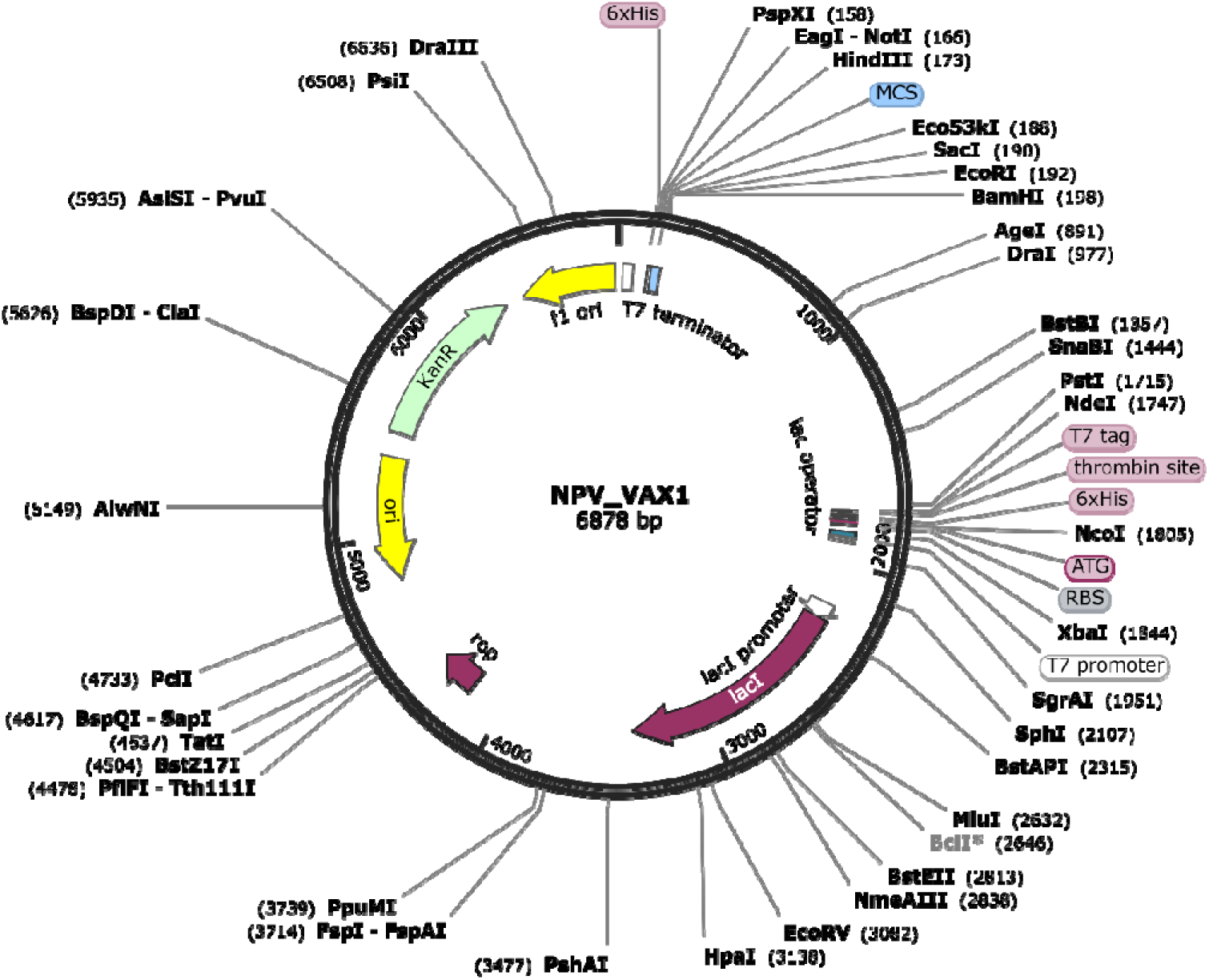
In silico cloning of the codon-optimized VC033_V6_BD_L03 construct into the pET-28a(+) expression vector. Circular plasmid map of the 6,878-bp recombinant pET-28a(+)-VC033_V6_BD_L03 construct generated using SnapGene. The optimized 1,539-bp vaccine coding sequence was inserted using the NdeI and BamHI restriction sites. Major vector features, including the T7 promoter, ribosome-binding site (RBS), His-tag regions, lacI, kanamycin-resistance marker (KanR), and origin of replication, are indicated.

## Discussion

Nipah virus remains an important target for vaccine development because of its recurrent zoonotic emergence, severe clinical outcomes, and the continuing absence of a licensed human vaccine. In the present study, a consensus-guided pan-proteome immunoinformatics strategy was used to prioritize a multi-epitope vaccine candidate from the complete nine-protein NiV repertoire. The workflow incorporated 901 protein sequences, generated consensus-derived sequence fragments, applied multi-stage epitope screening, and subsequently evaluated 60 vaccine constructs produced from alternative epitope-selection strategies, adjuvants, and sequence architectures. Eleven constructs satisfied the final construct-level selection criteria, from which VC033_V6_BD_L03 was prioritized for structural, receptor-interaction, molecular-dynamics, immune-simulation, and expression-oriented analyses. The principal strength of this approach is therefore the combination of broad proteome representation with sequence conservation and systematic construct-level optimization rather than reliance on a single viral antigen or a single arbitrarily assembled multi-epitope sequence.

The use of consensus-derived sequences represents an important distinction from most previous NiV immunoinformatics studies. Consensus-antigen strategies have already been explored experimentally for NiV: Choi et al. (2022) [15] generated consensus F and G sequences from multiple NiV isolates, while Lu et al. (2023) [16] generated an F-protein consensus sequence from GenBank sequences and demonstrated protection against both Malaysian and Bangladesh NiV strains in vaccinated Syrian hamsters. These studies provide experimental support for the broader concept of incorporating sequence diversity into antigen design, although both were focused on surface glycoproteins rather than the complete proteome. Among computational multi-epitope studies, Srivastava et al. (2023) [13] screened epitopes from all nine NiV proteins, but their use of “consensus” referred primarily to overlap among different epitope classes and prediction methods rather than generation of protein-level consensus sequences. Banico et al. (2024) [14] likewise incorporated all nine proteins and sequence variants, but identified conserved regions using alignment followed by Protein Variability Server analysis rather than constructing protein-specific consensus sequences. Soltan et al. (2021) [10] began with the complete nine-protein proteome but selected F, G, V, and W for subsequent epitope-based vaccine construction. Thus, to our knowledge, protein-level consensus generation across all nine NiV proteins followed by systematic screening of multiple multi-epitope vaccine architectures has not previously been reported for NiV vaccine design.

The multi-epitope strategy also allowed several complementary selection properties to be considered simultaneously. CTL, HTL, and B-cell candidates were screened for antigenicity, allergenicity, toxicity, sequence conservation, class-specific immune-response characteristics, and human-proteome similarity before vaccine assembly. HLA representation was retained as a major selection feature because broad HLA recognition is particularly relevant for an epitope-based vaccine intended for genetically diverse populations [39]. Population-coverage analysis showed combined predicted MHC-I/MHC-II coverage of 99.73–99.97% worldwide and 99.92–100% in South Asia among the 11 shortlisted constructs. VC033 itself reached 99.78% worldwide and 99.96% in South Asia. These values are consistent with, and in some cases exceed, those reported in previous NiV multi-epitope studies. Srivastava et al. (2023) [13] reported worldwide combined coverage of 97.88%, whereas Sharma et al. (2025) [12] reported 99.99% worldwide and 99.99% in South Asia. An additional strength of the present workflow was the evaluation of 60 alternative vaccine constructs, permitting epitope-selection strategy, adjuvant choice, linker arrangement, solubility, antigenicity, stability, hydropathicity, and related sequence characteristics to contribute to construct prioritization. The emergence of 11 hBD-3-containing constructs from this screening and subsequent selection of the V6/L03 design illustrates the value of evaluating vaccine architecture rather than assuming that one epitope arrangement will produce the most suitable complete construct.

Structural analysis provided further support for prioritization of VC033. The refined model showed 93.9% of residues in PROCHECK favored regions, an ERRAT quality factor of 91.192, and a standalone MolProbity score of 0.99. Structure-based B-cell analysis further identified 26 ElliPro discontinuous epitopes, of which 20 overlapped at least one DiscoTope-3.0-positive residue, and 10 showed at least 50% residue-level overlap. The agreement between ElliPro and DiscoTope-3.0 for multiple regions provides additional support for the presence of potentially antibody-accessible surfaces within the modeled construct [50,51]. Together with the favorable sequence-level properties of VC033, these analyses supported its progression to receptor-interaction screening.

A particularly notable result emerged from the TLR3 docking analysis. VC033_V6_BD_L03 produced a ClusPro model-0 score of −1624.8 against TLR3 structure 1ZIW, the most negative value among the constructs evaluated in the present study. This score also compares favorably with previously published NiV multi-epitope vaccine studies using ClusPro. Soltan et al. (2021) [10] reported a lowest ClusPro score of −1263.9 for a NiV vaccine–TLR3 complex using the same TLR3 structure, 1ZIW. Sharma et al. (2025) [12], also using ClusPro 2.0 and TLR3 1ZIW, reported scores of −1284.8 and −1222.6 for their NiV_1 and NiV_2 constructs, respectively. Other reported NiV ClusPro–TLR3 results include values down to −1089.1 in an integrated reverse-vaccinology study and −674.4 for a 2024 NiV multi-epitope construct docked to TLR3 7C76; a more recent computational mRNA vaccine study reported −1149.5 against TLR3. To our knowledge, the −1624.8 score obtained for VC033 is therefore the most negative ClusPro score reported to date for a NiV multi-epitope vaccine candidate docked with TLR3. The comparison with Soltan et al. (2021) [10] and Sharma et al. (2025) [12] is particularly informative because both the docking platform and TLR3 structure were the same. Independent analysis using HDOCK further ranked the VC033–1ZIW complex first, with a docking score of −314.68 and confidence score of 0.9642, strengthening its selection for subsequent dynamic evaluation.

The 100-ns molecular-dynamics simulation showed marked conformational adjustment during the early trajectory followed by a lower-variability regime after approximately 35–40 ns. During the final 50 ns, backbone RMSD averaged 1.250 ± 0.033 nm and Rg averaged 3.431 ± 0.017 nm. Intermolecular hydrogen bonding became more persistent during the later trajectory, with an average of 2.67 ± 1.14 contacts and at least one interchain hydrogen bond present in approximately 98.4% of sampled frames. PCA showed that PC1 and PC2 together accounted for 68.03% of positional variance, while the first six components accounted for 86.2%. MM/GBSA analysis yielded a mean interaction-energy estimate of −20.52 ± 15.42 kcal/mol, with favorable electrostatic and van der Waals contributions forming the major negative components. These results extend the docking analysis by showing persistence of receptor–construct contacts during the sampled trajectory. Previous NiV vaccine studies have also employed molecular dynamics or normal-mode approaches following TLR docking, including 100-ns simulations by Srivastava et al. (2023) [13] and Sharma et al. (2025) [12], whereas Soltan et al. (2021) [10] used iMODS normal-mode analysis.

The predicted immune-response profile was also consistent with the intended multi-epitope design. The primary two-dose C-ImmSim simulation produced B-cell and helper T-cell responses with persistent memory populations, a transient cytotoxic T-cell response, increased immunoglobulin production after the second administration, and prominent IFN-γ and IL-2 responses. An exploratory third administration on day 56 generated an additional simulated cellular, antibody, and cytokine response. Similar C-ImmSim approaches have been employed in other NiV multi-epitope studies, including those of Kumar et al. (2024) [11], Banico et al. (2024) [14], and Sharma et al. (2025) [12]. Expression-oriented analysis complemented these immunological predictions: the optimized 1,539-bp VC033 coding sequence showed a CAI of 0.75, GC content of approximately 50.3%, no internal NdeI or BamHI sites, and was successfully incorporated in silico into pET-28a(+), generating a 6,878-bp recombinant plasmid. Together, the immune-simulation and expression analyses provide a practical basis for moving the prioritized sequence toward recombinant production and experimental immunological evaluation.

The study has several limitations that should guide subsequent work. The entire vaccine-selection framework is computational, and the predicted epitope, population-coverage, antigenicity, allergenicity, toxicity, and immune-simulation outputs require experimental confirmation. Confidence in the modeled full-length structure was limited by the low ColabFold pLDDT values and modest QMEANDisCo scores, which also affect structure-dependent analyses such as conformational B-cell prediction and receptor docking. The production topology contained no explicit counterions, retained a net system charge of +23, and was represented by a single 100-ns trajectory. TLR3 docking should also be experimentally examined if receptor engagement is proposed as part of the construct’s immunological activity. These limitations define clear priorities for subsequent validation rather than altering the computational ranking achieved in the present workflow.

Overall, the present study prioritizes VC033_V6_BD_L03 as a lead NiV multi-epitope vaccine candidate through a workflow combining consensus-derived sequence representation of all nine viral proteins, extensive epitope screening, systematic comparison of 60 construct designs, broad predicted HLA coverage, structural and conformational-epitope analysis, receptor docking, molecular dynamics, immune simulation, and expression optimization. Particularly notable is the ClusPro score of −1624.8 against TLR3 1ZIW, which is more negative than previously reported NiV multi-epitope vaccine–TLR3 ClusPro scores identified in the published literature. The next stage should focus on recombinant expression and solubility, peptide–HLA validation, antibody recognition, and cellular immunogenicity, followed by in vivo assessment of immune responses and protective efficacy.

