## Supplementary File 1 for "Development of a Pan-Proteome Multi-Epitope Vaccine Candidate Against Nipah Virus Using Reverse Vaccinology and Immunoinformatics Approaches"

| **Protein Name** | **Query Keyword** |
|---|---|
| Glycoprotein | Nipah henipavirus[Organism] AND (glycoprotein[Title] OR "attachment protein"[Title]) NOT (partial[Title] OR fragment[Title] OR chain[Title] OR mutant[Title] OR synthetic[Title] OR construct[Title]) AND ("570"[SLEN] : "630"[SLEN]) |
| Fusion protein | Nipah henipavirus[Organism] AND "fusion protein"[Title] NOT (partial[Title] OR fragment[Title] OR chain[Title] OR mutant[Title] OR synthetic[Title] OR construct[Title]) AND ("520"[SLEN] : "570"[SLEN]) |
| Nucleocapsid protein | Nipah henipavirus[Organism] AND nucleocapsid[Title] NOT (partial[Title] OR fragment[Title] OR chain[Title] OR mutant[Title] OR synthetic[Title] OR construct[Title]) AND ("500"[SLEN] : "560"[SLEN]) |
| Matrix_Protein | Nipah henipavirus[Organism] AND matrix[Title] NOT (partial[Title] OR fragment[Title] OR chain[Title] OR mutant[Title] OR synthetic[Title] OR construct[Title]) AND ("330"[SLEN] : "370"[SLEN]) |
| Large_Polymerase protein | Nipah henipavirus[Organism] AND (polymerase[Title] OR "large protein"[Title]) NOT (partial[Title] OR fragment[Title] OR chain[Title] OR mutant[Title] OR synthetic[Title] OR construct[Title]) AND ("2100"[SLEN] : "2300"[SLEN]) |
| Phosphoprotein | Nipah henipavirus[Organism] AND phosphoprotein[Title] NOT (partial[Title] OR fragment[Title] OR chain[Title] OR mutant[Title] OR synthetic[Title] OR construct[Title]) AND ("670"[SLEN] : "740"[SLEN]) |
| V protein | Nipah henipavirus[Organism] AND "V protein"[Title] NOT (partial[Title] OR fragment[Title] OR chain[Title] OR mutant[Title] OR synthetic[Title] OR construct[Title]) AND ("430"[SLEN] : "480"[SLEN]) |
| W protein | Nipah henipavirus[Organism] AND "W protein"[Title] NOT (partial[Title] OR fragment[Title] OR chain[Title] OR mutant[Title] OR synthetic[Title] OR construct[Title]) AND ("410"[SLEN] : "450"[SLEN]) |
| C protein | Nipah henipavirus[Organism] AND "C protein"[Title] NOT (partial[Title] OR fragment[Title] OR chain[Title] OR mutant[Title] OR synthetic[Title] OR construct[Title]) AND ("160"[SLEN] : "190"[SLEN]) |
