## Supplementary File 2 for "Development of a Pan-Proteome Multi-Epitope Vaccine Candidate Against Nipah Virus Using Reverse Vaccinology and Immunoinformatics Approaches"

| Panel | Alleles |
|---|---|
| MHC-I human 27 referenc e panel | HLA-A*01:01, HLA-A*02:01, HLA-A*02:03, HLA-A*02:06, HLA-A*03:01, HLA-A*11:01, HLA-A*23:01, HLA-A*24:02, HLA-A*26:01, HLA-A*30:01, HLA-A*30:02, HLA-A*31:01, HLA-A*32:01, HLA-A*33:01, HLA-A*68:01, HLA-A*68:02, HLA-B*07:02, HLA-B*08:01, HLA-B*15:01, HLA-B*35:01, HLA-B*40:01, HLA-B*44:02, HLA-B*44:03, HLA-B*51:01, HLA-B*53:01, HLA-B*57:01, HLA-B*58:01 |
| MHC-II human 27 referenc e panel | HLA-DRB1*01:01, HLA-DRB1*03:01, HLA-DRB1*04:01, HLA-DRB1*04:05, HLA-DRB1*07:01, HLA-DRB1*08:02, HLA-DRB1*09:01, HLA-DRB1*11:01, HLA-DRB1*12:01, HLA-DRB1*13:02, HLA-DRB1*15:01, HLA-DRB3*01:01, HLA-DRB3*02:02, HLA-DRB4*01:01, HLA-DRB5*01:01, HLA-DQA1*05:01/DQB1*02:01, HLA-DQA1*05:01/DQB1*03:01, HLA-DQA1*03:01/DQB1*03:02, HLA-DQA1*04:01/DQB1*04:02, HLA-DQA1*01:01/DQB1*05:01, HLA-DQA1*01:02/DQB1*06:02, HLA-DPA1*02:01/DPB1*01:01, HLA-DPA1*01:03/DPB1*02:01, HLA-DPA1*01:03/DPB1*04:01, HLA-DPA1*03:01/DPB1*04:02, HLA-DPA1*02:01/DPB1*05:01, HLA-DPA1*02:01/DPB1*14:01 |
