## Supplementary File 3 for "Development of a Pan-Proteome Multi-Epitope Vaccine Candidate Against Nipah Virus Using Reverse Vaccinology and Immunoinformatics Approaches"

| construct_version | version_name | version_description |
|---|---|---|
| V1 | Highest antigenicity per protein | Selects the highest VaxiJen-scoring B-cell, CTL, and HTL epitope for each protein where available; fills missing class slots by the same antigenicity rule. |
| V2 | B/HTL antigenicity plus CTL immunogenicity | B-cell and HTL use highest VaxiJen score per protein; CTL uses highest MHC-I immunogenicity score per protein; fills missing class slots by the same class rule. |
| V3 | Maximum HLA allele breadth | B-cell uses highest VaxiJen score; CTL uses highest HLA-I allele count; HTL uses highest HLA-II allele count; fills missing class slots by the same class rule. |
| V4 | Top antigenicity by epitope class | Selects the top 9 B-cell, top 9 CTL, and top 9 HTL epitopes by VaxiJen score globally by class, without enforcing one epitope per protein. |
| V5 | Balanced composite per protein | Uses normalized composite scores combining antigenicity, immune breadth or cytokine scores, evidence count, and conservancy as available for each class. |
| V6 | CTL-strength construct | Prioritizes CTL epitopes by a composite of MHC-I immunogenicity, HLA-I allele count, and antigenicity; B-cell and HTL use antigenicity. |
| V7 | HTL-strength construct | Prioritizes HTL epitopes by a composite of HLA-II allele count, antigenicity, IFN-gamma, IL-4, and IL-10 scores; CTL uses immunogenicity and B-cell uses antigenicity. |
| V8 | Population-coverage proxy by class | Selects top 9 epitopes globally by class, prioritizing HLA-I allele breadth for CTL, HLA-II allele breadth for HTL, and antigenicity for B-cell. |
| V9 | Low human-similarity preference | Keeps all BLASTp-eligible epitopes but ranks lower human similarity higher, using antigenicity or CTL immunogenicity as a supporting score. |
| V10 | Low-overlap diversity construct | Selects protein-balanced epitopes while preferring lower coordinate overlap between epitopes from the same protein. |
