## Supplementary File 12 for "Development of a Pan-Proteome Multi-Epitope Vaccine Candidate Against Nipah Virus Using Reverse Vaccinology and Immunoinformatics Approaches"

| **(A)** | **(B)** |
| --- | --- |
| 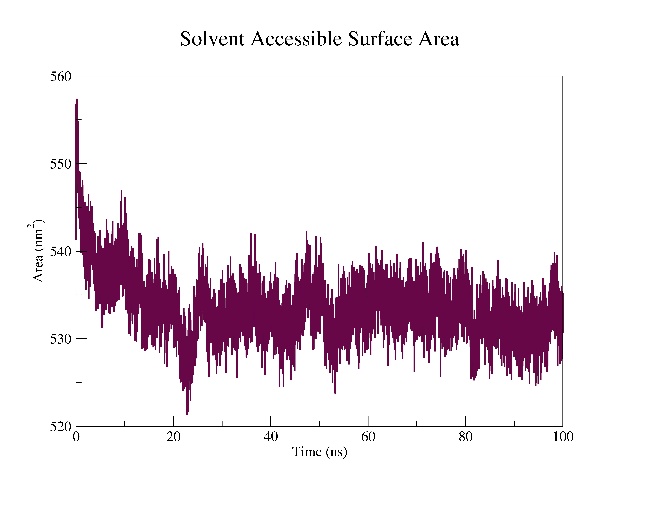 | 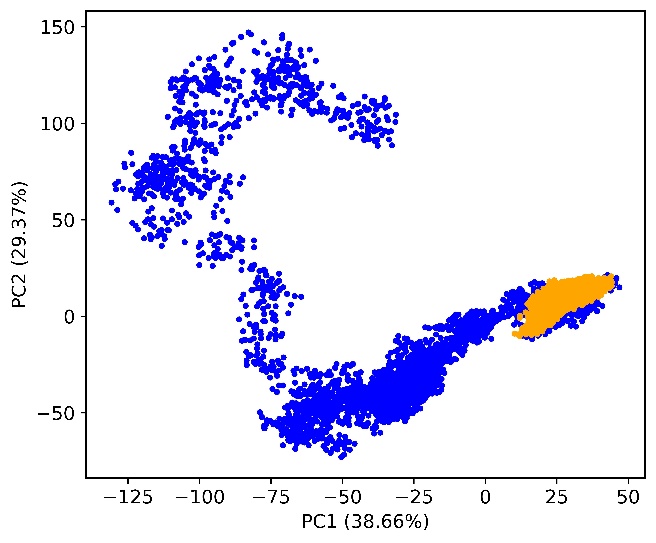 |
| **(C)** | **(D)** |
| 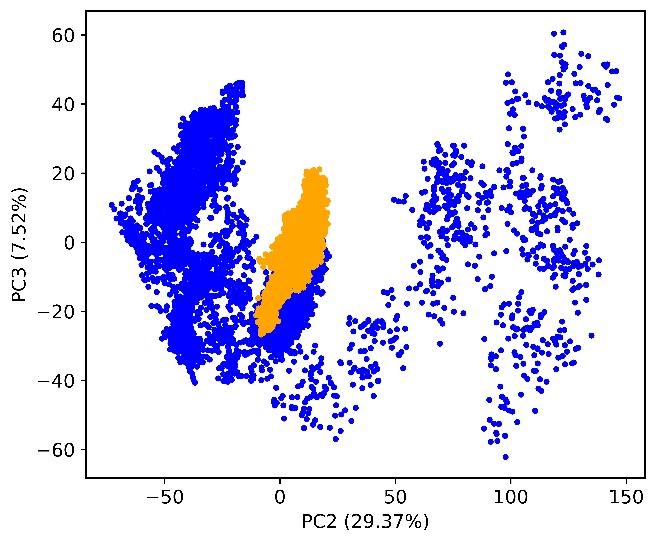 | 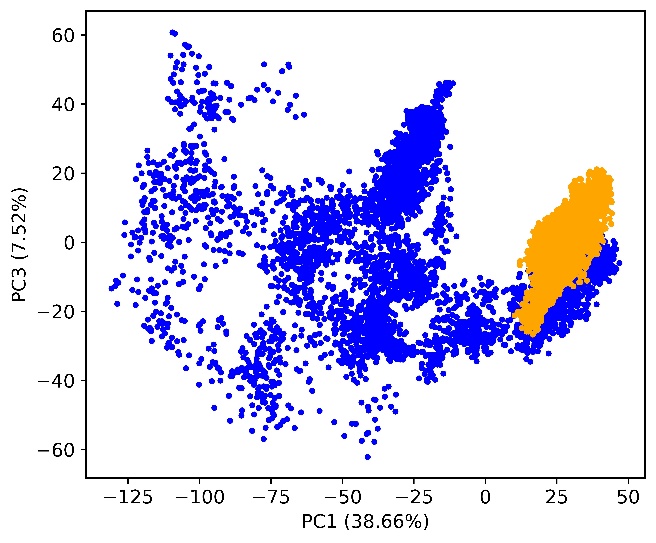 |
| **(E)** | **(F)** |
| 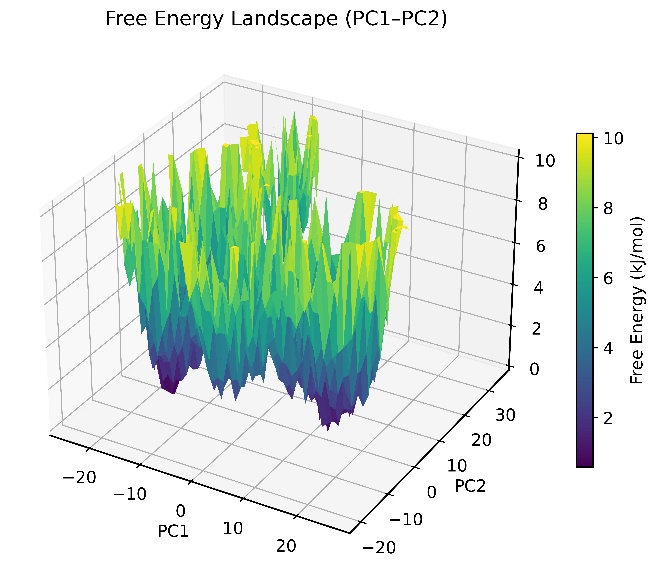 | 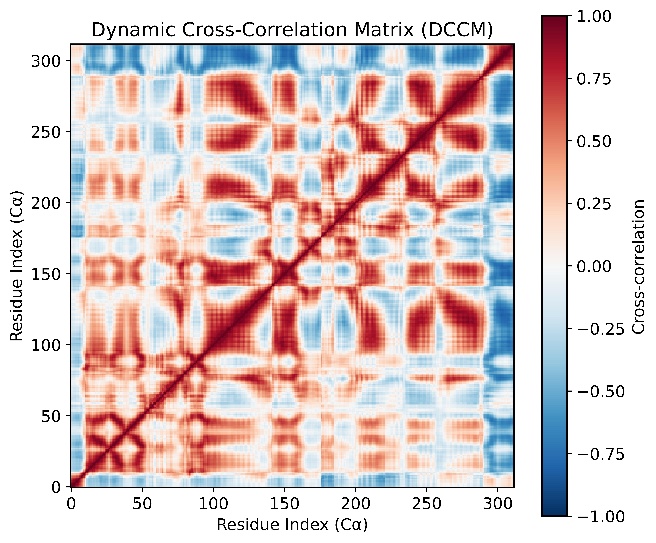 |

Figure 1. **Conformational dynamics and essential motion analysis.** **(A)** Solvent-accessible surface area (SASA) during the 100 ns simulation. **(B)** PCA projection of PC1–PC2. **(C)** PCA projection of PC2–PC3. **(D)** PCA projection of PC1–PC3. **(E)** Free-energy landscape (FEL) based on PC1 and PC2. **(F)** Dynamic cross-correlation matrix (DCCM) of residue motions.
