## Supplementary File 14 for "Development of a Pan-Proteome Multi-Epitope Vaccine Candidate Against Nipah Virus Using Reverse Vaccinology and Immunoinformatics Approaches"

### C-IMMSIM simulation results

August 16, 2026

**Abstract**

This document includes the plots relative to the simulation and the outcome of the epitope/peptide prediction used.

Produced by the C-IMMSIM Online server available at
`http://c-immsim.iac.rm.cnr.it (alias to http://kraken.iac.rm.cnr.it/C-IMMSIM)`

CITATIONS: For publication of results, please cite the following:

Nicolas Rapin, Ole Lund, Massimo Bernaschi, Filippo Castiglione. Computational Immunology Meets Bioinformatics: The Use of Prediction Tools for Molecular Binding in the Simulation of the Immune System. PLoS ONE 5(4): e9862
tt doi:10.1371/journal.pone.0009862, 2010

A retrospective validation

In-silico evaluation of adenoviral COVID-19 vaccination protocols: Assessment of immunological memory up to 6 months after the third dose. P. Stolfi, F. Castiglione, E. Mastrostefano, I. Di Biase, S. Di Biase, G. Palmieri, A. Prisco. Frontiers in Immunology, 13 (2022) doi: 10.3389/fimmu.2022.998262
`https://www.frontiersin.org/articles/10.3389/fimmu.2022.998262i`

An in vivo validation

Identification and validation of viral antigens sharing sequence and structural homology with tumor associated antigens (TAAs). C. Ragone, C. Manolio, B. Cavalluzzo, A. Petrizzo, A. Mauriello, M-L. Tornesello, F. M. Buonaguro, F. Castiglione, L. Vitagliano, M. Ruvo, M. Tagliamonte, L. Buonaguro. Journal for ImmunoTherapy of Cancer. 9:e002694 (2021)
`https://jitc.bmj.com/content/9/5/e002694`

Original C-IMMSIM model: `www.iac.cnr.it/~filippo/c-immsim`

GETTING HELP:
Scientific problems: Filippo Castiglione (filippo dot castiglione at cnr dot it)
Technical problems: Ilaria Gonnella (ilaria dot gonnella at cnr dot it)

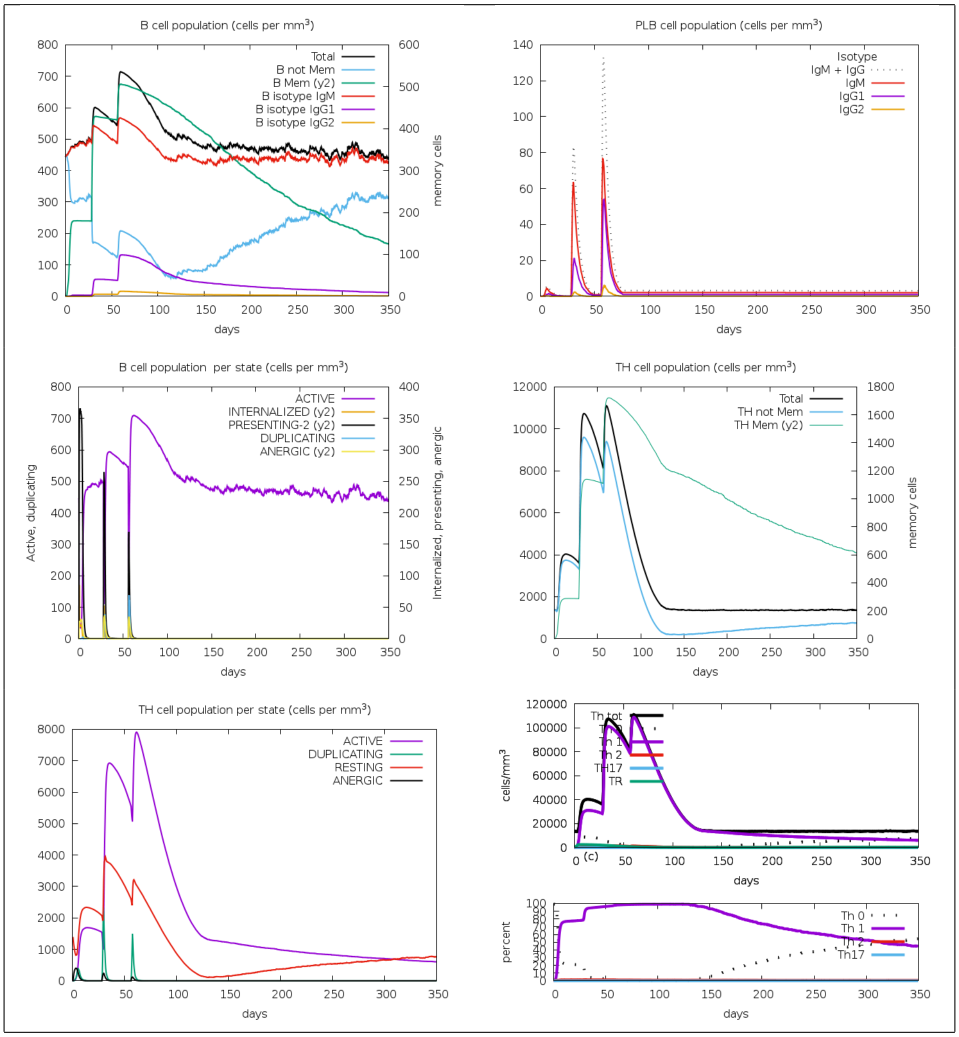


Figure 1: Cell counts shown. Legend: Act=active, Intern=internalized the Ag, Pres II = presenting on MHC II, Dup = in the mitotic cycle, Anergic = anergic, Resting = not active.

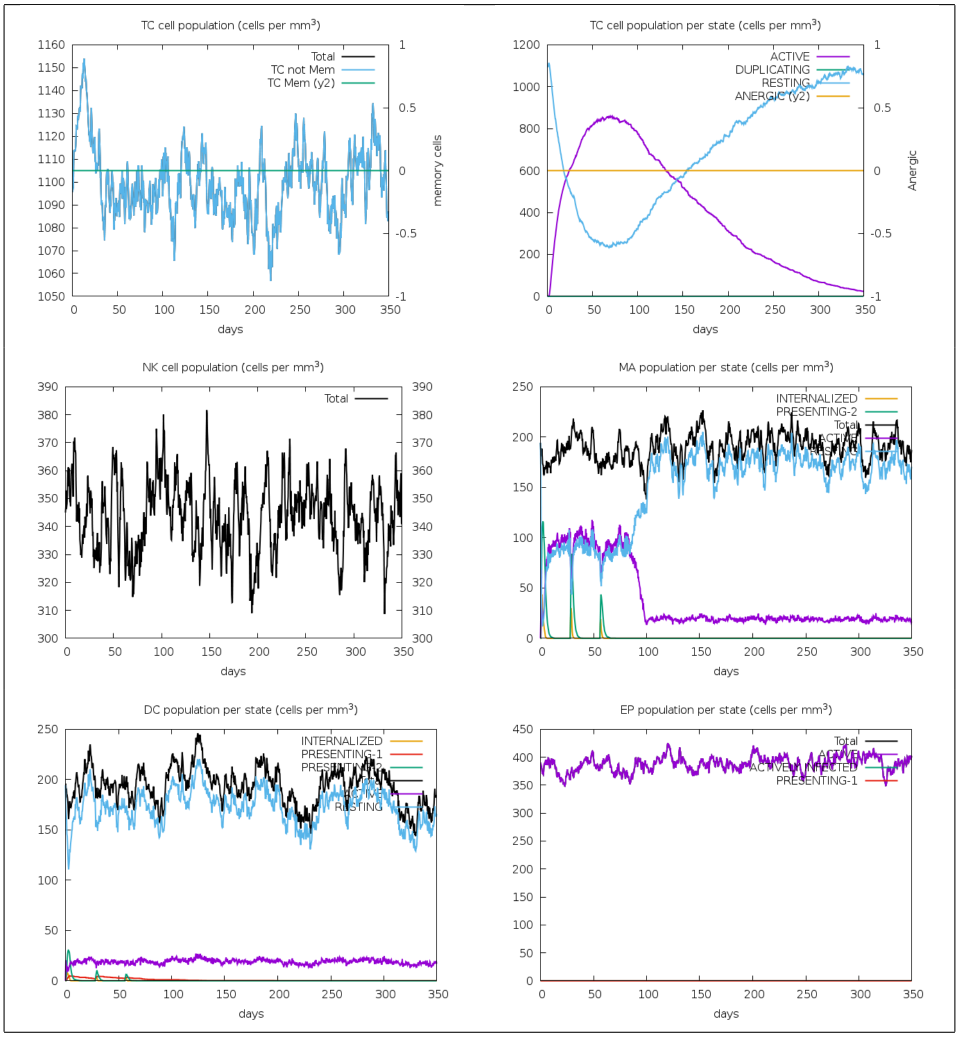


Figure 2: Legend: symbols as figure above.

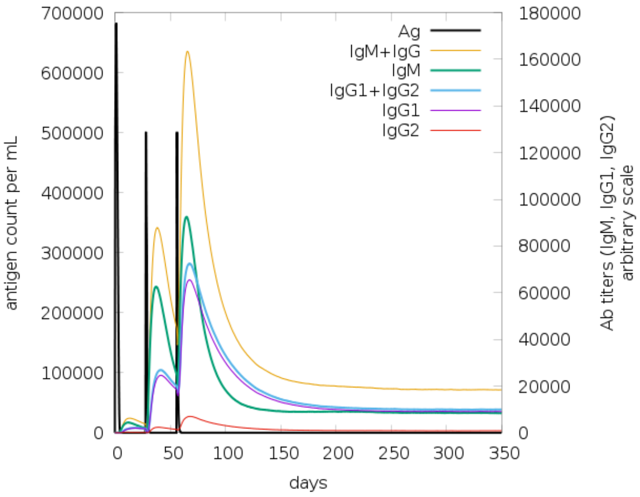


Figure 3: The virus, the immunoglobulins and the immunocomplexes.

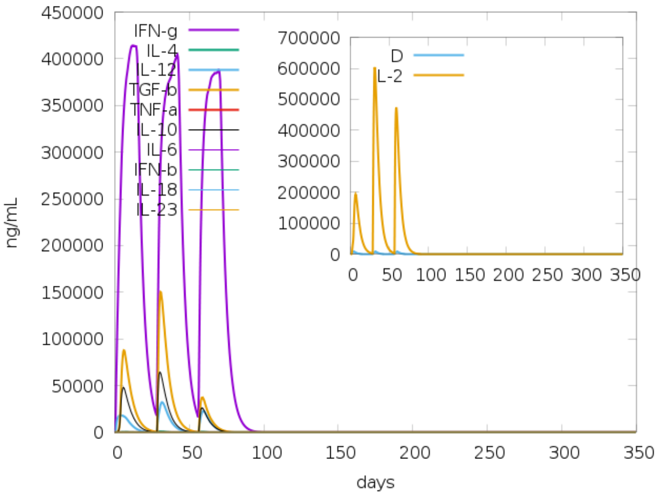


Figure 4: Concentration of cytokines and interleukins. Inset plot shows danger signal together with leukocyte growth factor IL-2.

---

```
Parker_B_scale_epitope_find
Use Parker's propensity scale, takes an antigen block as input,
and creates a list of residues that are possible epitopes.


----------------------------------
Antigen sequence file: /opt/lampp/htdocs/C-IMMSIM/Jobs/input/56582_20260813-200048_5_dOM4JM6NTUcn.FSA_1_001
----------------------------------


GIINTLQKYYCRVRGGRCAVLSCLPKEEQIGKCSTRGRKCCRRKKEAAAKLLTLFRRTKAAYISFIIVEKKAAYAFLIDR
INWAAYTIATIPFLFAAYALRIFFLSITKAAYTIRFGLETRAAYDQLEFEDEFAAYSSSEVIVGIAAYKSRGIPIKKHEY
GAEALERAGISCKQTELSLDLALSGPGPGFLLKNKIWCISLVEIGPGPGNIRRKVLILDFRSKLGPGPGQMGFQKNLCFS
LMDIGPGPGGRQSVTFKREMSISSGPGPGKNKIWCISLVEIYDTGPGPGKISCKQTELSLDLALGPGPGDKISCKQTELS
LDLAGPGPGDKSTEIILNPNLICIHEYGAEALERAGMASILLTLFRRTKKKYKKIGLITFISFIIVEKKRKKSSTITIPA
NIGLLGSKKKSIAITQKVHPNLPYKVKKAQMGFQKNLCFSLMDIKKDRSMGALNINRGYLEPKKFSFDNVKNFRDGSLTN
KKNTRDWAEGSDDIQLDPKKLNIIDFIQKNQKE

00000000000000000000000000011110011111111111111000000000000000000000000000000000
00000000000000000000000000000000000000000110000000000000100000000000000000000011
00000000000000000000000000000000000000000000000000000000000000000000000000000000
00000001111100000000000101111000000000000000000111000000000000101011000000000000
00000011111100000000000000000000000000000000000011100000000000000000111111000000
00000001100000000000000000000000000000000000010110000000000000000000000001001111
111101011111010000000000000001111
Epitopes of protein 0 ------------
1]       pos=27 len=4        EQIG
2]       pos=33 len=14        STRGRKCCRRKKEA
3]       pos=247 len=5        PGGRQ
4]       pos=265 len=4        PGPG
5]       pos=326 len=6        GPGDKS
6]       pos=388 len=6        KRKKSS
7]       pos=476 len=8        SLTNKKNT
8]       pos=487 len=5        AEGSD
9]       pos=509 len=4        NQKE
```

---

---

```
DoPeptideList_I:
        Given the antigen injected creates the list of peptides for all the
        NumAgProts proteins and for all i.e., 4 MHCI molecules

        Read class I peptide list from file? NO

=========================================================================

Allele: A0301
Pseudo sequence: KAAHVEQRKAQTRRV
Threshold: 8.185000
Max score: 23.064000


----------------------------------
Antigen sequence file: /opt/lampp/htdocs/C-IMMSIM/Jobs/input/56582_20260813-200048_5_dOM4JM6NTUcn.FSA_1_001
----------------------------------

GIINTLQKYYCRVRGGRCAVLSCLPKEEQIGKCSTRGRKCCRRKKEAAAKLLTLFRRTKAAYISFIIVEKKAAYAFLIDR
INWAAYTIATIPFLFAAYALRIFFLSITKAAYTIRFGLETRAAYDQLEFEDEFAAYSSSEVIVGIAAYKSRGIPIKKHEY
GAEALERAGISCKQTELSLDLALSGPGPGFLLKNKIWCISLVEIGPGPGNIRRKVLILDFRSKLGPGPGQMGFQKNLCFS
LMDIGPGPGGRQSVTFKREMSISSGPGPGKNKIWCISLVEIYDTGPGPGKISCKQTELSLDLALGPGPGDKISCKQTELS
```

```
LDLAGPGPGDKSTEIILNPNLICIHEYGAEALERAGMASILLTLFRRTKKKYKKIGLITFISFIIVEKKRKKSSTITIPA
NIGLLGSKKKSIAITQKVHPNLPYKVKKAQMGFQKNLCFSLMDIKKDRSMGALNINRGYLEPKKFSFDNVKNFRDGSLTN
KKNTRDWAEGSDDIQLDPKKLNIIDFIQKNQKE

Epitopes of protein 0 ------------
0]        pos=   1 score=0.000814 unnormalised=0.1070000000        IINTLQKYY
1]        pos=  61 score=0.006849 unnormalised=0.9000000000        YISFIIVEK
2]        pos= 100 score=0.061645 unnormalised=8.1010000000        RIFFLSITK
3]        pos= 140 score=0.006103 unnormalised=0.8020000000        VIVGIAAYK
4]        pos= 148 score=0.018225 unnormalised=2.3950000000        KSRGIPIKK
5]        pos= 214 score=0.001004 unnormalised=0.1320000000        VLILDFRSK
6]        pos= 261 score=0.006628 unnormalised=0.8710000000        ISSGPGPGK
7]        pos= 361 score=0.026930 unnormalised=3.5390000000        LTLFRRTKK
8]        pos= 362 score=0.033574 unnormalised=4.4120000000        TLFRRTKKK
9]        pos= 379 score=0.005859 unnormalised=0.7700000000        FISFIIVEK
10]        pos= 416 score=0.034570 unnormalised=4.5430000000        KVHPNLPYK
11]        pos=  -1 score=0.797798 unnormalised=104.8410000000        non-binding event

==========================================================================

Allele: A1101
Pseudo sequence: KAAHAEQRNKAQTRV
Threshold: 10.683000
Max score: 24.578000


----------------------------------
Antigen sequence file: /opt/lampp/htdocs/C-IMMSIM/Jobs/input/56582_20260813-200048_5_dOM4JM6NTUcn.FSA_1_001
----------------------------------

GIINTLQKYYCRVRGGRCAVLSCLPKEEQIGKCSTRGRKCCRRKKEAAAKLLTLFRRTKAAYISFIIVEKKAAYAFLIDR
INWAAYTIATIPFLFAAYALRIFFLSITKAAYTIRFGLETRAAYDQLEFEDEFAAYSSSEVIVGIAAYKSRGIPIKKHEY
GAEALERAGISCKQTELSLDLALSGPGPGFLLKNKIWCISLVEIGPGPGNIRRKVLILDFRSKLGPGPGQMGFQKNLCFS
LMDIGPGPGGRQSVTFKREMSISSGPGPGKNKIWCISLVEIYDTGPGPGKISCKQTELSLDLALGPGPGDKISCKQTELS
LDLAGPGPGDKSTEIILNPNLICIHEYGAEALERAGMASILLTLFRRTKKKYKKIGLITFISFIIVEKKRKKSSTITIPA
NIGLLGSKKKSIAITQKVHPNLPYKVKKAQMGFQKNLCFSLMDIKKDRSMGALNINRGYLEPKKFSFDNVKNFRDGSLTN
KKNTRDWAEGSDDIQLDPKKLNIIDFIQKNQKE

Epitopes of protein 0 ------------
0]        pos= 100 score=0.032219 unnormalised=3.5800000000        RIFFLSITK
1]        pos= 140 score=0.001719 unnormalised=0.1910000000        VIVGIAAYK
2]        pos= 361 score=0.011862 unnormalised=1.3180000000        LTLFRRTKK
3]        pos= 416 score=0.010656 unnormalised=1.1840000000        KVHPNLPYK
4]        pos=  -1 score=0.943544 unnormalised=104.8410000000        non-binding event

==========================================================================

Allele: B1501
Pseudo sequence: KAAREEQQIKTQTRE
Threshold: 9.167400
Max score: 25.365000


----------------------------------
Antigen sequence file: /opt/lampp/htdocs/C-IMMSIM/Jobs/input/56582_20260813-200048_5_dOM4JM6NTUcn.FSA_1_001
----------------------------------

GIINTLQKYYCRVRGGRCAVLSCLPKEEQIGKCSTRGRKCCRRKKEAAAKLLTLFRRTKAAYISFIIVEKKAAYAFLIDR
INWAAYTIATIPFLFAAYALRIFFLSITKAAYTIRFGLETRAAYDQLEFEDEFAAYSSSEVIVGIAAYKSRGIPIKKHEY
GAEALERAGISCKQTELSLDLALSGPGPGFLLKNKIWCISLVEIGPGPGNIRRKVLILDFRSKLGPGPGQMGFQKNLCFS
LMDIGPGPGGRQSVTFKREMSISSGPGPGKNKIWCISLVEIYDTGPGPGKISCKQTELSLDLALGPGPGDKISCKQTELS
LDLAGPGPGDKSTEIILNPNLICIHEYGAEALERAGMASILLTLFRRTKKKYKKIGLITFISFIIVEKKRKKSSTITIPA
NIGLLGSKKKSIAITQKVHPNLPYKVKKAQMGFQKNLCFSLMDIKKDRSMGALNINRGYLEPKKFSFDNVKNFRDGSLTN
KKNTRDWAEGSDDIQLDPKKLNIIDFIQKNQKE
```

```
Epitopes of protein 0 ------------
0]       pos=  46 score=0.004199 unnormalised=0.4636000000        AAAKLLTLF
1]       pos=  56 score=0.019082 unnormalised=2.1066000000        RTKAAYISF
2]       pos=  94 score=0.006446 unnormalised=0.7116000000        FAAYALRIF
3]       pos= 464 score=0.020604 unnormalised=2.2746000000        FSFDNVKNF
4]       pos=  -1 score=0.949669 unnormalised=104.8410000000        non-binding event

=========================================================================

Allele: B3501
Pseudo sequence: KAARVEQQIKAQTRE
Threshold: 9.376400
Max score: 27.550000


----------------------------------
Antigen sequence file: /opt/lampp/htdocs/C-IMMSIM/Jobs/input/56582_20260813-200048_5_dOM4JM6NTUcn.FSA_1_001
----------------------------------

GIINTLQKYYCRVRGGRCAVLSCLPKEEQIGKCSTRGRKCCRRKKEAAAKLLTLFRRTKAAYISFIIVEKKAAYAFLIDR
INWAAYTIATIPFLFAAYALRIFFLSITKAAYTIRFGLETRAAYDQLEFEDEFAAYSSSEVIVGIAAYKSRGIPIKKHEY
GAEALERAGISCKQTELSLDLALSGPGPGFLLKNKIWCISLVEIGPGPGNIRRKVLILDFRSKLGPGPGQMGFQKNLCFS
LMDIGPGPGGRQSVTFKREMSISSGPGPGKNKIWCISLVEIYDTGPGPGKISCKQTELSLDLALGPGPGDKISCKQTELS
LDLAGPGPGDKSTEIILNPNLICIHEYGAEALERAGMASILLTLFRRTKKKYKKIGLITFISFIIVEKKRKKSSTITIPA
NIGLLGSKKKSIAITQKVHPNLPYKVKKAQMGFQKNLCFSLMDIKKDRSMGALNINRGYLEPKKFSFDNVKNFRDGSLTN
KKNTRDWAEGSDDIQLDPKKLNIIDFIQKNQKE

Epitopes of protein 0 ------------
0]       pos=  94 score=0.003902 unnormalised=0.4286000000        FAAYALRIF
1]       pos= 115 score=0.009110 unnormalised=1.0006000000        FGLETRAAY
2]       pos= 139 score=0.021101 unnormalised=2.3176000000        EVIVGIAAY
3]       pos= 356 score=0.011359 unnormalised=1.2476000000        MASILLTLF
4]       pos=  -1 score=0.954528 unnormalised=104.8410000000        non-binding event
```

---

---

```
DoPeptideList_II:
        Given the antigen injected creates the list of peptides for all the
        NumAgProts proteins and for all i.e., 2 MHCII molecules

        Read class II peptide list from file? NO

=========================================================================

Allele: DRB1_0301
Pseudo sequence: RVTLGRDAYW
Threshold: 2.237910
Max score: 30.179000


----------------------------------
Antigen sequence file: /opt/lampp/htdocs/C-IMMSIM/Jobs/input/56582_20260813-200048_5_dOM4JM6NTUcn.FSA_1_001
----------------------------------

GIINTLQKYYCRVRGGRCAVLSCLPKEEQIGKCSTRGRKCCRRKKEAAAKLLTLFRRTKAAYISFIIVEKKAAYAFLIDR
INWAAYTIATIPFLFAAYALRIFFLSITKAAYTIRFGLETRAAYDQLEFEDEFAAYSSSEVIVGIAAYKSRGIPIKKHEY
GAEALERAGISCKQTELSLDLALSGPGPGFLLKNKIWCISLVEIGPGPGNIRRKVLILDFRSKLGPGPGQMGFQKNLCFS
LMDIGPGPGGRQSVTFKREMSISSGPGPGKNKIWCISLVEIYDTGPGPGKISCKQTELSLDLALGPGPGDKISCKQTELS
LDLAGPGPGDKSTEIILNPNLICIHEYGAEALERAGMASILLTLFRRTKKKYKKIGLITFISFIIVEKKRKKSSTITIPA
NIGLLGSKKKSIAITQKVHPNLPYKVKKAQMGFQKNLCFSLMDIKKDRSMGALNINRGYLEPKKFSFDNVKNFRDGSLTN
KKNTRDWAEGSDDIQLDPKKLNIIDFIQKNQKE
```

```
Epitopes of protein 0 ------------
0]       pos=   1 score=0.008994 unnormalised=1.5140900000        IINTLQKYY
1]       pos=  50 score=0.006113 unnormalised=1.0290900000        LLTLFRRTK
2]       pos=  53 score=0.029161 unnormalised=4.9090900000        LFRRTKAAY
3]       pos=  65 score=0.064666 unnormalised=10.8860900000        IIVEKKAAY
4]       pos=  75 score=0.019360 unnormalised=3.2590900000        FLIDRINWA
5]       pos=  93 score=0.004931 unnormalised=0.8300900000        LFAAYALRI
6]       pos= 102 score=0.000832 unnormalised=0.1400900000        FFLSITKAA
7]       pos= 103 score=0.027041 unnormalised=4.5520900000        FLSITKAAY
8]       pos= 115 score=0.046228 unnormalised=7.7820900000        FGLETRAAY
9]       pos= 176 score=0.025092 unnormalised=4.2240900000        LSLDLALSG
10]       pos= 215 score=0.091712 unnormalised=15.4390900000        LILDFRSKL
11]       pos= 255 score=0.020239 unnormalised=3.4070900000        FKREMSISS
12]       pos= 297 score=0.008091 unnormalised=1.3620900000        LSLDLALGP
13]       pos= 360 score=0.006113 unnormalised=1.0290900000        LLTLFRRTK
14]       pos= 363 score=0.011602 unnormalised=1.9530900000        LFRRTKKKY
15]       pos= 383 score=0.022057 unnormalised=3.7130900000        IIVEKKRKK
16]       pos= 384 score=0.010960 unnormalised=1.8450900000        IVEKKRKKS
17]       pos= 403 score=0.027243 unnormalised=4.5860900000        LLGSKKKSI
18]       pos= 404 score=0.004307 unnormalised=0.7250900000        LGSKKKSIA
19]       pos= 439 score=0.033135 unnormalised=5.5780900000        SLMDIKKDR
20]       pos= 443 score=0.053647 unnormalised=9.0310900000        IKKDRSMGA
21]       pos= 464 score=0.013336 unnormalised=2.2450900000        FSFDNVKNF
22]       pos= 476 score=0.019520 unnormalised=3.2860900000        SLTNKKNTR
23]       pos= 493 score=0.052566 unnormalised=8.8490900000        IQLDPKKLN
24]       pos=  -1 score=0.393054 unnormalised=66.1680000000        non-binding event

========================================================================

Allele: DRB1_0401
Pseudo sequence: KFAAHVEQRKAQTRV
Threshold: 2.842330
Max score: 29.818000


---------------------------------
Antigen sequence file: /opt/lampp/htdocs/C-IMMSIM/Jobs/input/56582_20260813-200048_5_dOM4JM6NTUcn.FSA_1_001
---------------------------------

GIINTLQKYYCRVRGGRCAVLSCLPKEEQIGKCSTRGRKCCRRKKEAAAKLLTLFRRTKAAYISFIIVEKKAAYAFLIDR
INWAAYTIATIPFLFAAYALRIFFLSITKAAYTIRFGLETRAAYDQLEFEDEFAAYSSSEVIVGIAAYKSRGIPIKKHEY
GAEALERAGISCKQTELSLDLALSGPGPGFLLKNKIWCISLVEIGPGPGNIRRKVLILDFRSKLGPGPGQMGFQKNLCFS
LMDIGPGPGGRQSVTFKREMSISSGPGPGKNKIWCISLVEIYDTGPGPGKISCKQTELSLDLALGPGPGDKISCKQTELS
LDLAGPGPGDKSTEIILNPNLICIHEYGAEALERAGMASILLTLFRRTKKKYKKIGLITFISFIIVEKKRKKSSTITIPA
NIGLLGSKKKSIAITQKVHPNLPYKVKKAQMGFQKNLCFSLMDIKKDRSMGALNINRGYLEPKKFSFDNVKNFRDGSLTN
KKNTRDWAEGSDDIQLDPKKLNIIDFIQKNQKE

Epitopes of protein 0 ------------
0]       pos=  54 score=0.038848 unnormalised=4.6796700000        FRRTKAAYI
1]       pos=  75 score=0.023465 unnormalised=2.8266700000        FLIDRINWA
2]       pos=  93 score=0.011561 unnormalised=1.3926700000        LFAAYALRI
3]       pos= 102 score=0.030065 unnormalised=3.6216700000        FFLSITKAA
4]       pos= 104 score=0.004015 unnormalised=0.4836700000        LSITKAAYT
5]       pos= 135 score=0.006066 unnormalised=0.7306700000        YSSSEVIVG
6]       pos= 141 score=0.014234 unnormalised=1.7146700000        IVGIAAYKS
7]       pos= 169 score=0.003907 unnormalised=0.4706700000        ISCKQTELS
8]       pos= 176 score=0.057045 unnormalised=6.8716700000        LSLDLALSG
9]       pos= 219 score=0.017024 unnormalised=2.0506700000        FRSKLGPGP
10]       pos= 255 score=0.133020 unnormalised=16.0236700000        FKREMSISS
11]       pos= 290 score=0.003907 unnormalised=0.4706700000        ISCKQTELS
12]       pos= 297 score=0.035038 unnormalised=4.2206700000        LSLDLALGP
13]       pos= 311 score=0.003907 unnormalised=0.4706700000        ISCKQTELS
14]       pos= 443 score=0.068609 unnormalised=8.2646700000        IKKDRSMGA
15]       pos=  -1 score=0.549290 unnormalised=66.1680000000        non-binding event
```
