## Supplementary File 15 for "Development of a Pan-Proteome Multi-Epitope Vaccine Candidate Against Nipah Virus Using Reverse Vaccinology and Immunoinformatics Approaches"

| Web server / resource | Verified URL |
| --- | --- |
| NCBI Protein database | https://www.ncbi.nlm.nih.gov/protein/ |
| CD-HIT Suite web server | http://cd-hit.org |
| Clustal Omega - EMBL-EBI Job Dispatcher | https://www.ebi.ac.uk/jdispatcher/msa/clustalo |
| EMBOSS Cons - EMBL-EBI Job Dispatcher | https://www.ebi.ac.uk/jdispatcher/msa/emboss_cons |
| IEDB T Cell Prediction, Class I (TC1) | https://nextgen-tools.iedb.org/pipeline?tool=tc1 |
| NetMHCpan 4.1 - DTU Health Tech | https://services.healthtech.dtu.dk/services/NetMHCpan-4.1/ |
| IEDB T Cell Prediction, Class II (TC2) | https://nextgen-tools.iedb.org/pipeline?tool=tc2 |
| NetMHCIIpan 4.3 - DTU Health Tech | https://services.healthtech.dtu.dk/services/NetMHCIIpan-4.3/ |
| IEDB B-cell Sequence Prediction | https://nextgen-tools.iedb.org/pipeline?tool=bcell_sequence |
| BepiPred 3.0 - DTU Health Tech | https://services.healthtech.dtu.dk/services/BepiPred-3.0/ |
| ABCpred | https://webs.iiitd.edu.in/raghava/abcpred/ |
| VaxiJen v2.0 | https://www.ddg-pharmfac.net/vaxijen/VaxiJen/VaxiJen.html |
| AllerTOP v2.0 | http://www.ddg-pharmfac.net/AllerTOP |
| ToxinPred | https://webs.iiitd.edu.in/raghava/toxinpred/ |
| IEDB Epitope Conservancy Analysis | https://tools.iedb.org/conservancy/ |

| Web server / resource | Verified URL |
|---|---|
| IEDB Class I Immunogenicity | https://tools.iedb.org/immunogenicity/ |
| IFNepitope2 | https://webs.iiitd.edu.in/raghava/ifnepitope2/ |
| IL4pred | https://webs.iiitd.edu.in/raghava/il4pred/ |
| IL10pred | https://webs.iiitd.edu.in/raghava/il10pred/ |
| NCBI Protein BLAST (BLASTp) | https://blast.ncbi.nlm.nih.gov/Blast.cgi?PAGE=Proteins |
| ToxinPred2 | https://webs.iiitd.edu.in/raghava/toxinpred2/ |
| ExPASy ProtParam | https://web.expasy.org/protparam/ |
| SoluProt | https://loschmidt.chemi.muni.cz/soluprot/ |
| IEDB Population Coverage | https://tools.iedb.org/population/ |
| PSIPRED Workbench | https://bioinf.cs.ucl.ac.uk/psipred/ |
| NetSurfP-3.0 - DTU Health Tech | https://services.healthtech.dtu.dk/services/NetSurfP-3.0/ |
| ColabFold AlphaFold2_mmseqs2 - Google Colab | https://colab.research.google.com/github/sokrypton/ColabFold/blob/main/AlphaFold2.ipynb |
| GalaxyRefine - GalaxyWEB | https://galaxy.seoklab.org/refine |
| SAVES v6.1 Structure Validation Server | https://saves.mbi.ucla.edu/ |
| MolProbity | https://molprobity.biochem.duke.edu/ |
| QMEAN / QMEANDisCo | https://swissmodel.expasy.org/qmean/ |
| ProSA-web | https://prosa.services.came.sbg.ac.at/prosa.php |

| Web server / resource | Verified URL |
|---|---|
| IEDB B-cell Structure Prediction | https://nextgen-tools.iedb.org/pipeline?tool=bcell_structure |
| ClusPro 2.0 | https://cluspro.bu.edu/ |
| HDOCK | http://hdock.phys.hust.edu.cn/ |
| C-IMMSIM / C-ImmSim | https://kraken.iac.rm.cnr.it/C-IMMSIM/index.php |
| EMBOSS Backtranseq - EMBL-EBI Job Dispatcher | https://www.ebi.ac.uk/jdispatcher/st/emboss_backtranseq |
| GenSmart Codon Optimization Tool | https://www.genscript.com/tools/gensmart-codon-optimization |
| GenRCA Rare Codon Analysis Tool | https://www.genscript.com/tools/rare-codon-analysis |
| RNAfold - ViennaRNA Web Services | http://rna.tbi.univie.ac.at/cgi-bin/RNAWebSuite/RNAfold.cgi |
| SnapGene | https://www.snapgene.com/ |
